# Exploring the effect of Hydrocarbon Cross-linkers on the Structure and Binding of Stapled p53 Peptides

**DOI:** 10.1101/2024.10.01.616000

**Authors:** Asha Rani Choudhury, Vikram Gaikwad, Atanu Maity, Rajarshi Chakrabarti

## Abstract

Short-length peptides are used as therapeutics due to their high target specificity and low toxicity, for example, peptides designed for targeting the interaction between oncogenic protein p53 and E3 ubiquitin ligase MDM2. These peptide therapeutics form a class of successful inhibitors. To design such peptide-based inhibitors, stapling is one of the methods in which amino acid side chains are stitched together to get conformationally rigid peptides, ensuring effective binding to their partners. In the current work, we use computer simulations to investigate p53 peptides stapled with hydrocarbon chains of different lengths and positions of attachment to the peptide. We subsequently analyze their binding efficiency with MDM2. The introduction of stapling agents restricts the conformational dynamics of peptides, resulting in higher persistence of helicity. The efficiency of the stapling agents has also been verified imposing these stapled peptides to adverse conditions viz. thermal and chemical denaturation. In addition, the conformational exploration of peptides has been investigated using Temperature replica exchange molecular dynamics (T-REMD) simulations. From both the unbiased and T-REMD simulations, p53 with a long hydrocarbon cross-linker shows a more conformationally rigid structure having high helicity compared to other stapled peptides. The rigidity gained due to cross-linking reduces the entropy of the peptide in the free state and thereby facilitates the complexation process. From the binding studies, we have shown that the peptide having multiple short staples has shown a larger enthalpy change during binding free energy, resulting from its orientation and interactions of residues in the binding interface. On the other hand, a peptide with a single long stapling agent shows less entropic penalty than other systems. Our studies suggest a plausible rationale for the relation between the length and the position of attachment of cross-linkers to peptides and the binding free energies between the peptides and their target partners.

## 1. Introduction

Proteins are essential biomolecules serving in various biological functions such as enzyme catalysis, hormone regulation, signal transduction, and gene expression. Proteins interact with other biomolecules and the complex networks of these interactions characterize different steps involved in biomolecular processes. Protein-protein interaction is one of these key interactions. The protein-protein interaction is very much signified in terms of their specificity for binding, diversity, evolution, and potential to serve as therapeutics.^1^ Dysregulation of these sophisticated interactions results in malfunctioning of the biomolecular processes which leads to numerous diseases such as cancer, neurodegenerative disorders, and infections.^2^ In diseased cells, often some protein-protein interactions are over expressed or the natural binding partner of a protein is replaced by another protein. Designing therapeutics against a disease involves finding promising strategies to interfere into these altered protein-protein interactions and restore a healthy cellular environment. This is usually done by designing molecules that can replace one partner in the protein-protein interaction.^3^ Those therapeutics are broadly classified into 3 categories, small molecules,^4^ peptides^5^, and biologics, based on their size, molecular weight, cell permeability, and target selectivity. Amongst these therapeutics, peptides lie in between small molecules and biologics. Peptide therapeutics have proven a potential method to combat diseases and many of the designed peptides are now in clinical trials.^6^ Basically, peptide therapeutics are short chains of amino acids that can bind to the specific region of its targeting binding partner and modulate various biological processes. Some of them are orally bioavailable, cell-permeable, and target specific.^7^ Peptide therapeutics are designed to mimic the binding of naturally occurring proteins which are highly efficient and selective. The promising applications of peptide therapeutics are found in the treatment of various diseases such as cancer, cardiovascular diseases, and metabolic disorders. In tumor genesis, the protein-protein interaction of p53-MDM2 plays a vital role. Inhibition of this interaction is a key challenge in the design of p53 peptide therapeutics.

The tumor suppressor p53 plays a vital role in apoptosis, DNA repair, cell-cycle arrest, etc.; for which, it is also called the ‘Guardian of the genome’.^8^ It activates the cellular pathway for apoptosis upon DNA damage and cellular stress. In normal cellular conditions, the level of p53 is controlled by E3 ubiquitin ligase murine double minute (MDM2). In order to prevent the accumulation of p53 in normal cell growth and division, MDM2 binds to p53 and propagates ubiquitination along with proteasomal degradation.^9^ Thus, MDM2 is also known as a negative regulator of p53. But in the tumor cells, MDM2 is often overexpressed leading to suppression of the p53 activity, which leads to uncontrolled cell growth.^10,11^ This E3 ubiquitin ligase MDM2 binds with p53 by interacting through its N-terminal transactivation domain (NTD). This NTD of p53 has an alpha-helical domain that resides in the hydrophobic cleft of MDM2 by anchoring its 3 hydrophobic residues Phe19, Trp23, and Leu26.^12^ So to restore the p53 tumor suppressing activity, it is crucial to inhibit the interaction between MDM2 and p53, various small molecules,^13,14^ and peptide inhibitors,^12,15^ have been designed for this purpose. Amongst these therapeutics, peptide inhibitors show stronger binding with MDM2 due to their specificity. However, one of the challenges in the case of peptide inhibitors is the low membrane permeability and proteolytic degradation.

To overcome this challenge in the p53-MDM2 interaction, designing an alpha-helical part of p53 that interacts with MDM2 by maintaining the spatial positions of interacting residues can resolve the problem.^16^ Designing a small peptide with a high degree of helicity while maintaining the crucial amino acid positions is challenging. Over the past several decades, researchers have developed various non-covalent and covalent strategies to induce helicity in these shorter peptides so that they can act as peptide-based drugs.^17^ One of the recognized covalent methods is known as “stapling” or “cross-linking”.^18^ In this type of chemical modification, the side chains of suitably positioned amino acids are replaced by different types of chemical groups and linked together to form a cross-linker attached peptide termed as “stapled peptide” which successfully maintains its backbone rigidity and thus reduces the entropic penalty. Additionally, this strategy has also been proven successful for membrane permeability.^19^ Recently, stapled peptides have been extensively studied and found efficient as peptide therapeutics.^6,20,21^ The designing of these stapled peptides can be regulated by two determining factors (i) chemical nature of the cross-linker such as alkenyl, aryl, triazole, disulfide, lactam.^22,23,24,25^ and (ii) length and position of cross-linker as along the peptide sequence, such as i-i+3 (connecting the i^th^ and the i+3^rd^ residues), i-i+4, i-i+7, i-i+11.^26^ The hydrocarbon cross-linkers are dominantly used for stapling purposes while the optimized positions for staple are i-i+4 and i-i+7. The efficacy of hydrocarbon cross-linkers has been demonstrated by different developed models.^27^ Also depending upon the length of the peptide inhibitor and the position of the staple, more than one stapling agent can be used to get a desirable affinity and target specificity.^21^ Other than p53-MDM2 interaction, there are other examples where stapled peptides have been successfully used as therapeutic agents such as the BCL pathway,^28^ NOTCH pathway^29^, and in targeting HIV^30^.

In past years, remarkable progress has been made in the development of stapled peptides and comparison among them has been done based on their kinetic and thermodynamic properties.^21,31,32^ These studies are mostly based on the conformational change of p53 in a bound state with MDM2. However, the introduction of a stapling agent into a peptide can change its conformation in the free state which eventually will affect its interaction with the target partner. This change in conformational dynamics of p53 in the free state involves a transition between helical and non-helical conformations which affects the entropic penalty of p53 binding with MDM2. Along with this, the residue level interaction of MDM2-p53 has a crucial role in their binding studies. In the case of p53-MDM2, an i-i+7 stapled peptide has been reported, and it shows effective binding at the MDM2 hydrophobic interface.^33^ Considering this approach, in our previous work, we designed a new aromatic staple that reduced the entropic cost even more than the i-i+7 aliphatic stapled peptide upon binding.^34^

In this work, we keep the chemical nature of the cross-linker identical and change the length and position of attachment to the peptide. We are curious to know its effect on the conformational dynamics of free peptides and the binding thermodynamics of p53-MDM2. Here, we have considered different lengths of hydrocarbon cross-linkers such as a single i-i+4 stapling agent, a single i-i+7 stapling agent, and a double i-i+4 aliphatic stapling agent for p53. We extensively studied the conformation of peptides cross-linked by the aforementioned stapling agents in an aqueous solution and also in denaturing conditions such as high temperature 330K and chemical denaturant 8M urea. We used the Temperature-Replica Exchange molecular dynamics simulation^35^ to investigate temperature dependent conformational exploration of modeled stapled p53 peptides. We compared the binding free energy of all these peptides with MDM2 using MM/GBSA calculations.^36^ The outcome of this study is to detect the potential stapled peptide therapeutic and its effective binding using computer simulations.

## 2. Method and simulation details

### 2.1 Modelling of stapled peptide

The structure of the helical domain of p53 bound with its natural binding partner E3 ubiquitin ligase MDM2 is considered the system of interest (PDB ID: 1YCR). This p53 (sequence: ETFSDLWKLLPEN) is a 13-residue stretch of human tumor suppressor p53 and has been used in some earlier studies for recapitulating various aspects of MDM2-p53 interaction.^34,37^

The human p53 protein is a long chain of 393 amino acids comprised of four functional domains. Out of which the amino-terminal transcriptional domain or NTD contains the transcriptional factor of the p53 (amino acids 1-42) region. Several proteins bind to the transcription activation domain of p53 and activate the expression of various target genes and MDM2 is one of them. The NTD of p53 interacting with MDM2, is unstructured to a greater extent. But its interaction with the target protein induces a conformational change in its structure and leads to an increase in the binding affinity which regulates various biological processes. Hence, maintaining the secondary structure of these 13 residues stretch p53 (termed as p53 in the rest of the manuscript) will be helpful for its binding with negative regulator MDM2.

In this work, we have used a stapling approach where the side chains of two amino acids of the wild-type p53 are joined together to form a stapled peptide.^18,20^ In this stapling approach, the stapling positions are selected without any disruption of hydrophobic contact of p53 peptide with its negative regulator MDM2 which is formed by 3 hydrophobic residues Phe3, Trp7, and Leu10. Considering all these criteria, aliphatic cross-linkers have been used in this study differing in their length and positions of attachments to p53.

p53 with a single i-i+4 staple is designed by replacing Ser4 and Lys8 of p53 and termed as p53^i-i+4^ in the rest of the article. Similarly, p53 with an i-i+7 aliphatic staple is obtained from crystal structure 3V3B^33^ where the aliphatic chain is attached at positions Ser4, and Pro11 and is denoted as p53^i-i+7^. Since a single i-i+4 is not able to span the overall length of peptide and also to probe the effectiveness of a single i-i+7 and double i-i+4 stapling agent, we have designed a double stapled p53 where two i-i+4 aliphatic chains are considered as cross-linkers between the two pairs of residues by replacing amino acid pairs such as THR2 - LEU6 and LYS8 – GLU12 and termed as p53^double^. All these aforementioned stapled peptides along with the wild-type version (represented in Figure 1) have been simulated in free state for a comparison study of their conformational behavior and their binding thermodynamics with targeting partner MDM2 has been investigated by simulating those in complex with MDM2. The details of these peptides and their complexes are listed in Table 1 with the simulation length and environment details.

**Figure 1:**
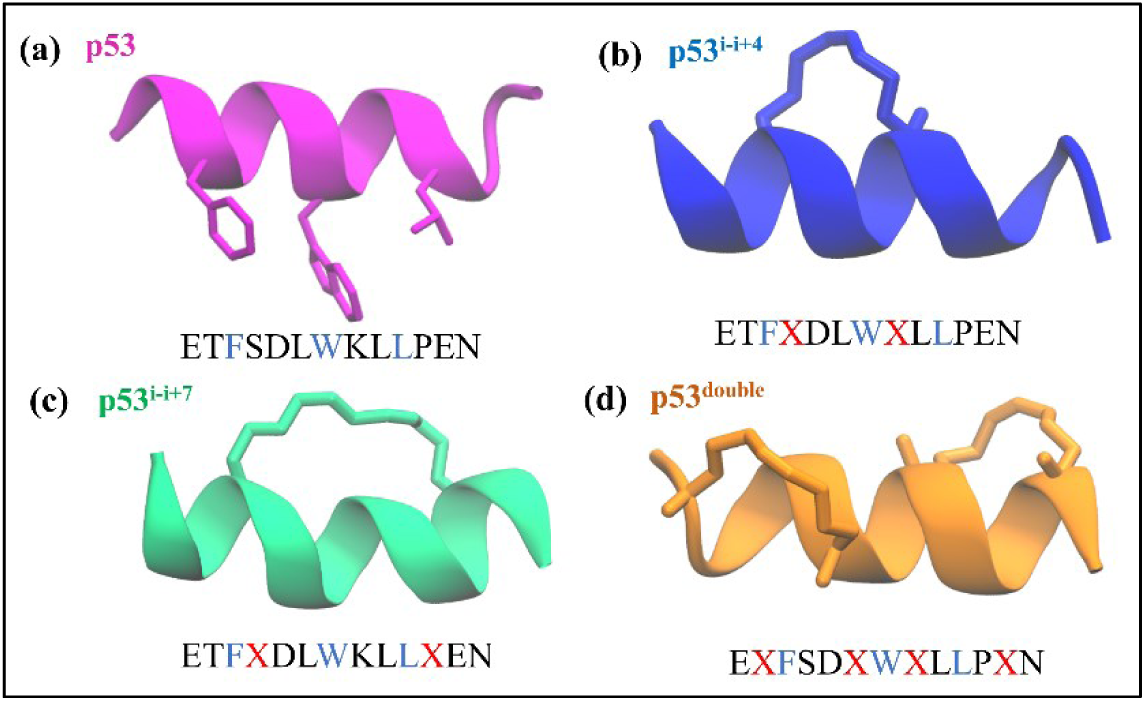
Sequences and structures of the four peptides - a) p53, b) p53^i-i+4^, c) p53^i-i+7^, and d) p53^double^. The stapling positions in all these peptides have been marked with X in the sequence and the anchoring residues are colored in blue. The peptide structures are shown in cartoon representations and the interacting hydrophobic residues F3, W7, and L10 for p53 and the stapling agents for p53^i-i+4^, p53^i-i+7^, and p53^double^ are shown in stick representations with their respective color.

**Table 1:** Details of the systems and simulation lengths for different systems. The system identifier defined in the following table is maintained throughout the manuscript.

| Sl No. | System Identifier | Protein | Peptide | Temp. (K) | Conc. of urea (M) | Simulation time (μs) |
| --- | --- | --- | --- | --- | --- | --- |
| 1 | p53 | - | p53 | 310 | - | 3 x 1 |
| 2 | p53_330K | - | p53 | 330 | - | 3 x 1 |
| 3 | p53_urea | - | p53 | 310 | 8 | 3 x 1 |
| 4 | p53<sup>i-i+4</sup> | - | p53 with a single i-i+4 aliphatic staple | 310 | - | 3 x 1 |
| 5 | p53<sup>i-i+4</sup>_330K | - | p53 with a single i-i+4 aliphatic staple | 330 | - | 3 x 1 |
| 6 | p53<sup>i-i+4</sup>_urea | - | p53 with a single i-i+4 aliphatic staple | 310 | 8 | 3 x 1 |
| 7 | p53<sup>i-i+7</sup> | - | p53 with a single i-i+7 aliphatic staple | 310 | - | 3 x 1 |
| 8 | p53<sup>i-i+7</sup>_330K | - | p53 with a single i-i+7 aliphatic staple | 330 | - | 3 x 1 |
| 9 | p53<sup>i-i+7</sup>_urea | - | p53 with a single i-i+7 aliphatic staple | 310 | 8 | 3 x 1 |
| 10 | p53<sup>double</sup> | - | p53 with double i-i+4 aliphatic staple | 310 | - | 3 x 1 |
| 11 | p53<sup>double</sup>_330K | - | p53 with double i-i+4 aliphatic staple | 310 | - | 3 x 1 |
| 12 | p53<sup>double</sup>_urea | - | p53 with double i-i+4 aliphatic staple | 310 | - | 3 x 1 |
| 13 | MDM2 | MDM2 | - | 310 | - | 3 x 1 |
| 14 | MDM2 + p53 | MDM2 | p53 | 310 | - | 3 x 1 |
| 15 | MDM2 + p53<sup>i-i+4</sup> | MDM2 | p53 with a single i-i+4 aliphatic staple | 310 | - | 3 x 1 |
| 16 | MDM2+p53<sup>i-i+7</sup> | MDM2 | p53 with a single i-i+7 aliphatic staple | 310 | - | 3 x 1 |
| 17 | MDM2+p53<sup>double</sup> | MDM2 | p53 with double i-i+4 aliphatic staple | 310 | - | 3 x 1 |
| | | | | | | 51 μs |

### 2.2 Simulation Details

The CHARMM36^38^ forcefield parameters were employed to design the stapled peptide and a combination of CHARMM parameters of proteins and small molecules (CGENFF)^39^ were used to model those stapling agents. For designing an aliphatic cross-linker at the position of the targeted residue, first, the amino acid side chains were replaced by a lysine group then ɛ-NH2 groups of lysine were replaced by an aliphatic chain having an olefinic bond, and the suitable length, and the end of the two side chains were joined together. This strategy has been successfully used to model stapled peptides in other studies as well.^40,41^ All of the peptides were capped at N-terminal and C-terminal by acetyl and amide groups respectively to avoid interaction of the bare terminal charges with the rest of the system.

In this work, we performed multiple all-atom unbiased simulations for free p53 peptides and their stapled variations in the free state and complex with the target partner MDM2. Equilibrium molecular dynamics simulations often sample the conformations near global minima region so to explore other regions and to check the efficiency of stapling agents on peptide stability different denaturing conditions were used such as (i) simulation at high temperature 330K which is slightly higher than the melting temperature of p53^42^ and (ii) simulation in the presence of 8M urea. These two conditions are known to be standard for studying the denaturation of proteins and short peptides such as p53.^43,44^On the other hand, to explore the larger conformational space of the designed peptides we have used temperature replica exchange molecular dynamics (T-REMD)^35^ simulation as one of the enhanced sampling approaches.

Finally, to probe the binding of these peptides with their targeting partner MDM2, we have performed multiple unbiased simulations for the three complex structures; MDM2+p53^wt^, MDM2+p53^i-i+7^, MDM2+p53^double^, and MDM2. The first two complex structures were generated as per their crystal structures 1YCR and 3V3B respectively while MDM2+p53^double^ is modeled by replacing p53^i-i+7^ with p53^double^ in its complex with MDM2. Simulation details of all these systems are provided in Table 1.

Each of the systems is neutralized by adding the required numbers of K^+^ and Cl^−^ ions and solvated in a cubic water box made of TIP3P water,^45^ where the size of the box is determined by maintaining a distance of at least 1 nm between the water box edge and peptide atoms to satisfy the periodic boundary condition. For the removal of initial steric clashes, 5000-step energy minimization is performed for each system using the steepest descent method^46^ and applying position restraint on heavy atoms. Subsequently, a 500 ps position restrained equilibration in the NVT ensemble is performed using V-rescale thermostat^47^ for equilibrating each system at 310K to avoid void formation in the box followed by a 20ns unrestrained equilibration is performed at isothermal-isobaric (NPT) ensemble to attain a steady pressure of 1 atm considering a pressure relaxation of 1ps using Parrinello-Rahman barostat.^48^ Finally, the production runs for 1000ns with a time step of 2 fs were performed for each system in the NPT ensemble. All the simulations are performed in GROMACS.^49^ Short-ranged Lennard– Jones interactions are calculated using the minimum image convention.^50^ For estimating non-bonding interactions including electrostatic as well as van der Waals interactions, a spherical cut-off distance of 1 nm is chosen. Periodic boundary conditions have been used in all three directions to remove edge effects. SHAKE algorithm^51^ is applied to constrain bonds involving the hydrogen atoms. Long-range electrostatic interactions are calculated using the particle mesh Ewald (PME) method.^52^ The coordinates from the simulation are saved at a frequency of 2 ps for analyses. To extract different structural properties and for visualization, in-built modules of GROMACS,^53^ VMD,^54^ PLUMED^55^ are used.

### 2.3 Replica exchange molecular dynamics simulation

The temperature replica exchange molecular dynamics (T-REMD)^35^ is one enhanced sampling technique for the exploration of the complex conformational energy landscape of biomolecules. In this method, multiple copies of the system are simulated in parallel, each at a different temperature and periodically exchanging their configurations between adjacent replicas. This exchange is accepted with a probability defined by the Metropolis criterion as,

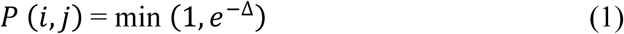

Where Δ = (*β*_*i*_ − *β*_*j*_)(*U*(*R*_*i*_) − *U*(*R*_*j*_))

*P* (*i*, *j*) is the acceptance probability between adjacent replicas i and j

*U*(*R*_*i*_) and *U*(*R*_*j*_) – Potential energy of replica i and j respectively

β = 1/*k_B_T* (*k*_*B*_ is Boltzmann constant and T is the temperature in Kelvin)

The temperature values of replicas were obtained using the temperature generator replica web server^56^ within the temperature range of 303-500K. After that 50ns production run has been done per replica with a time step of 2 fs and an exchange between adjacent replicas was done for every 100 steps, giving a total of 2.5 μs to 3.5 μs production simulation time for each system. Following this, the coordinates were saved for every 1 ps over the total production run for each replica. These simulations have been run by using the mpi module of the GROMACS 2020.6 package.^53^

**Table 2:** T-REMD simulation details are given in the following table.

| Sl. No. | System Identifier | Peptide | T-REMD Simulation Details |  |  |
| --- | --- | --- | --- | --- | --- |
|  |  |  | Number of Replicas | Temperature Range (K) | Production Time (ns) |
| 1 | p53 | p53 with no staple | 70 | 303 -500 | 50 x 70 |
| 2 | p53<sup>i-i+4</sup> | p53 with a single i-i+4 aliphatic staple | 51 | 303 - 500 | 50 x 51 |
| 3 | p53<sup>i-i+7</sup> | p53 with a single i-i+7 aliphatic staple | 68 | 303 - 500 | 50 x 68 |
| 4 | p53<sup>double</sup> | p53 with double i-i+4 aliphatic staple | 51 | 303 - 500 | 50 x 51 |
| | | | | | 12 μs |

A good exchange between replicas can be interpreted from the random walk of a trajectory in all temperature space. Here, the desired exchange acceptance ratio is given as 25% (Table ST1). Furthermore, the reasonable overlap of potential energies for the first 10 replicas shows a good exchange of conformations. This allows good sampling at all temperatures, with high-temperature simulations crossing the energy barriers and low-temperature simulations to explore local free energy minima. The trajectories derived from T-REMD are analyzed further to check the conformational exploration of peptides and their helical efficiency with respect to temperature. Along with this, clustering has also been done to get an idea about the representative conformations of populated states. These analyses were calculated by using Gromacs^53^ and PLUMED^55^ and some in-house scripts.

### 2.4 Binding free energy calculation

In this work, we have used the most popular and computationally less expensive method MM-GBSA (Molecular mechanics with generalized Born and surface area)^36^ method to calculate the relative free binding free energy between p53 peptides and MDM2. This method has been already used in protein-protein interaction studies to determine free energy and also to find the residue level interaction in their binding.^57^ A summary of the above method can be explained as,

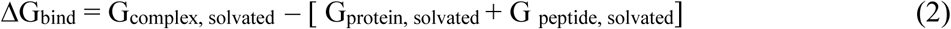

where ΔG_solvated_ can be represented as,

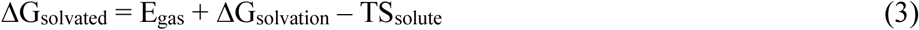

which can be further written as -

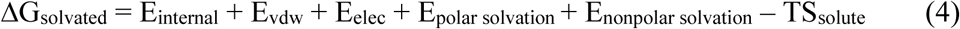

The first three terms of equation (4) describe the molecular mechanical (MM) gas-phase energy which arises due to molecular motion such as bond vibration, angle bending, dihedral rotation, and van der Waals and electrostatic interactions respectively. The fourth and fifth terms represent the solvation free energy calculated considering implicit environments in two separate parts, i.e., polar, and non-polar components. The polar solvation energy is estimated considering GB (Generalized Born) solvent model and the non-polar component is assumed to be proportional to the solvent-accessible surface area (SASA).^58,59^ The last term is solute entropy, where T represents the absolute temperature and S is the entropy of the solute, whereas the solvation entropy is already incorporated in solvation energy terms. The solute entropy is calculated by considering quasi-harmonic approximation.^60^ In short, the sum of the first five terms is considered as enthalpy, H although it includes the solvation entropy term and simply the binding free energy can be written as,

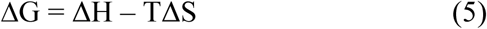

In practice, all the simulations of the free and complex state are performed in explicit water to get the conformational microstates of the solute, and then the free energy of solvation of these conformations is estimated in an implicit environment using generalized Born (GB) together with the surface-area term. In this work, we have followed multiple trajectory protocols (MTP)^61^ and performed three independent simulations for MDM2+p53 complex, MDM2, and p53 systems and their stapled versions. The enthalpy and its decomposition are calculated by gmx_MMGBSA.^62^ These calculations are averaged over 25,000 conformations for each system on the last 500ns simulations. The free energy values are summarised in Table 3 and a detail of the energy terms is provided in the supporting information table (Table ST2 to ST5). This method has been performed earlier in p53-MDM2 interaction to explain their interactions.^34,63^

**Table 3:** Free energies of MDM2-p53 binding and component wise contributions to overall binding free energy. All the energy values are in kcalmol^−1^.

| Energy Components | MDM2 + p53 | MDM2+ p53<sup>i-i+4</sup> | MDM2 + p53<sup>i-i+7</sup> | MDM2 + p53<sup>double</sup> |
| --- | --- | --- | --- | --- |
| ΔBOND | 3.49 ± 4.22 | -1.81 ± 0.43 | 5.39 ± 3.86 | -2.67 ± 0.05 |
| ΔANGLE | 2.26 ± 1.48 | -2.55 ± 0.16 | -1.61 ± 2.59 | -6.06 ± 2.52 |
| ΔDIHED | 1.48 ± 4.23 | 9.87 ± 4.13 | 5.70 ± 2.00 | 11.88 ± 5.96 |
| ΔUB | 0.11 ± 0.15 | -0.85 ± 0.10 | 0.26 ± 0.18 | -0.92 ± 0.10 |
| ΔIMP | 0.47 ± 0.26 | 0.01 ± 0.31 | 0.24 ± 0.20 | 0.02 ± 0.19 |
| ΔCMAP | 3.55 ± 1.54 | 5.06 ± 2.73 | 2.33 ± 0.40 | 1.73 ± 6.16 |
| ΔVDWAALS | -71.57 ± 8.96 | -76.60 ± 1.87 | -70.76 ± 3.68 | -71.69 ± 2.55 |
| Δ1-4 VDW | 2.02 ± 2.02 | 3.78 ± 0.31 | 3.12 ± 0.94 | 2.73 ± 1.42 |
| ΔEEL | -294.72 ± 50.05 | -340.21 ± 85.59 | -253.88 ± 17.72 | -255.61 ± 13.66 |
| Δ1-4 EEL | -1.50 ± 12.02 | 9.66 ± 3.38 | 16.59 ± 5.16 | -1.71 ± 12.92 |
| ΔG<sub>GAS</sub> | -354.39 ± 37.62 | -393.65 ± 82.67 | -292.63 ± 20.86 | -322.30 ± 23.59 |
| ΔEGB | 315.98 ± 39.76 | 337.87 ± 74.08 | 260.89 ± 13.37 | 261.18 ± 23.85 |
| ΔESURF | -10.03 ± 0.84 | -9.12 ± 1.20 | -9.05 ± 0.32 | -8.99 ± 0.27 |
| ΔG<sub>SOLV</sub> | 305.95 ± 38.92 | 328.75 ± 72.89 | 251.84 ± 13.39 | 252.19 ± 23.78 |
| Δ Enthalpy(ΔH) | -48.43 ± 1.48 | -64.89 ± 9.96 | -40.79 ± 7.49 | -70.11 ± 1.42 |
| -TΔS | 40.21 | 44.74 | 22.22 | 47.55 |
| ΔΔG | -8.22 | -20.14 | -18.56 | -22.56 |

## 3. Results and Discussion

The goal of the study is to determine how and to what extent the introduction of hydrocarbon cross-linkers of different lengths at various positions in the peptide sequence modifies the equilibrium conformational population of the peptides in the aqueous solution. These configurational dynamics have also been verified by using an enhanced sampling method, temperature replica exchange molecular dynamics (T-REMD) simulations. Also, the binding free energy of stapled peptides with MDM2 has been computed and a detailed analysis of per residue energy contribution to the binding of p53 with MDM2 is done.

### Unbiased Molecular dynamics simulations of free peptide

#### 3.1 Reduction of conformational flexibility

To investigate the conformational dynamics of peptides in the presence and absence of different stapling agents, a 2D - population based conformational landscape is constructed considering two important structural parameters of the peptide such as the Radius of gyration (Rg) and the backbone Root mean square deviation (RMSD) from 1 µs long unbiased simulation for each free peptides. These two parameters are conventionally used to describe the conformational space proficiently.^64^ The probability of occurrence of the points within the 0.1nm x 0.1nm grid is calculated from the R_g_ vs RMSD scatter plot using the following expression,

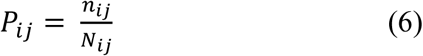

Where *n*_*ij*_ is the number of possible conformations within an area defined by i and i+Δi values of Rg and j and j+Δj values of RMSD. *N*_*ij*_ is the total number of conformations. The free energy is calculated using this probability by the following formula;

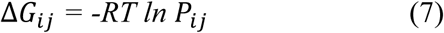

To explore the efficiency of the stapling agent to reduce the conformational flexibility of peptides, two denaturing conditions are imposed i.e. thermal denaturation and chemical denaturation. For thermal denaturation, the peptides were simulated at 330K, and for chemical denaturation, they were simulated in an 8M urea solution. The energy minimum points in the conformational energy landscape are the most populated regions during the simulation. The free energy plots (Figure 2) clearly show that the wild type p53 has a wider distribution in all three conditions compared to p53 stapled systems. Wild-type p53 covers a broad range of Rg and RMSD starting from lower values at the initial period of simulation to the higher values of these parameters when they undergo unfolding. But with the introduction of a stapling agent, the area of free energy surface is reduced in aqueous solution and denaturing conditions. However, for the p53^i-i+4^ peptide, the free energy surface is wider along the Rg axis. A plausible explanation for this behavior is due to the range of residues covered by the cross-linker is short in the above peptide. The energy minimum in the aqueous solution of p53^i-i+4^ is obtained at a value of 0.8nm Rg and 0.3nm RMSD. This RMSD value increases to higher values at 330K and in 8M urea solution while there is no such notable change in the minima along the Rg axis implying the overall shape of the peptide being maintained. A similar kind of behavior is also reflected in p53^i-i+7^ and p53^double^. Hence the introduction of cross-linkers helps to maintain the overall shape of the peptide though there are small deviations from their initial conformations. The free energy surfaces of p53^i-i+7^ and p53^double^ are more restricted compared to that of p53 and p53^i-i+4^. Here, for both of these cases, the populated basins occur at lower RMSD values, however for p53^double^, it is at a lower value of Rg than p53^i-i+7^. The p53^double^ samples a wider conformational landscape than p53^i-i+7^ in an aqueous solution. Similarly, in the case of high temperature, it samples more unfolded conformations than p53^i-i+7^, but in the presence of 8M urea, p53^double^ spans over a restricted conformational area. This implies less exposure of peptide backbone atoms to the urea molecules. This has been quantified in the radial distribution of urea around the peptide backbone atoms which exhibits reduced p53^double^ radial distribution (Figure S1). Overall, the reason for this restriction can be explained by the change in length and the number of cross-linkers over the peptide sequence.

**Figure 2:**
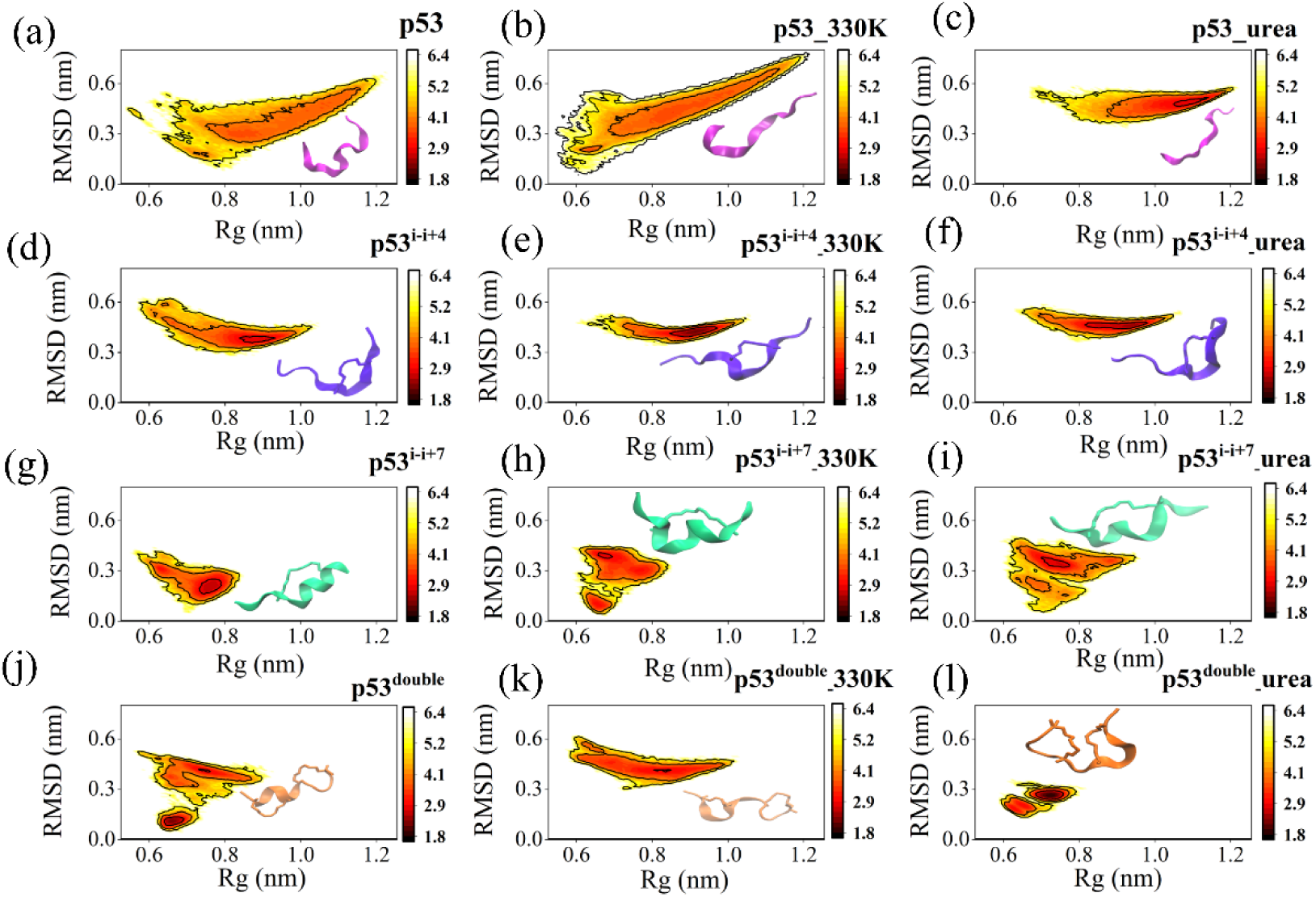
2D free energy profiles obtained from the R_g_ and RMSD values for the p53 peptide (a-c) and its stapled versions p53^i-i+4^ (d-f), p53^i-i+7^ (g-i), and p53^double^ (k-l) in the aqueous medium (first column) and different conditions (i.e. at high temperature, and in the presence of urea: 2^nd^ column and 3^rd^ column respectively) showing conformational rigidity in the presence of stapling agents. The increase in free energy change is depicted using a color gradient scale from dark brown to white color where the free energy values are in kcal mol^−1^. Hence, conformations represented by darker brown color are the major energy minimum implying those conformations are more populated than those conformations which are represented by light yellow color, and those populated structures are shown in insets. This plot is generated over the second half of 1µs unbiased simulation.

The acquired conformational rigidity of the peptides with the introduction of cross-linkers can be related to their residue level fluctuations which are quantified from the root mean square fluctuation plot (Figure S2). An observation of Figure S2 will give information about those residues that are contributing to the rigidity of the free energy landscape.

In addition to the conformational rigidity, the helical conformation of p53 is crucial for its interaction with MDM2. This helical propensity of the peptide is quantified using the formula implemented in PLUMED^55^ and the probability distributions of the helical values of the peptides in different conditions are given in supporting information (Figure S3).

From the Rg-RMSD free energy surface (Figure 2) and probability distribution of the alpha-helicity plot (Figure S3), it becomes evident that the cross-linkers are effective in reducing the conformational flexibility of peptides and in maintaining their helical nature in adverse conditions. Next, we focus on the effect of cross-linkers on the shape and structure of the free peptides and compare those properties based on population density. This type of analysis is crucial when one would try to analyze the variation in the length of the peptide with their respective helicity values. Thus, stapled peptide having a suitable cross-linker length and appropriate point of attachment can be identified as a potential alpha-helical inhibitor to its binding partner. The ultimate goal is to study the binding free energy between MDM2 and considered peptides, so further structural analysis of peptides is done in an aqueous condition.

#### 3.2 End-to-end distance and Helical fraction free energy surface

The native contact of the residues can be quantified by the end-to-end distance of a peptide. However, some earlier studies are showing conformationally populated landscapes over extended conformations where the α-helicity of peptides is still favored.^65^ To understand the above mechanism and change in structural properties of peptides with the attachment of cross-linker, a free energy surface is constructed using end-to-end distance and alpha-helicity as reaction coordinates as shown in Figure 3. This plot has been generated over the last 500ns of unbiased simulations.

**Figure 3:**
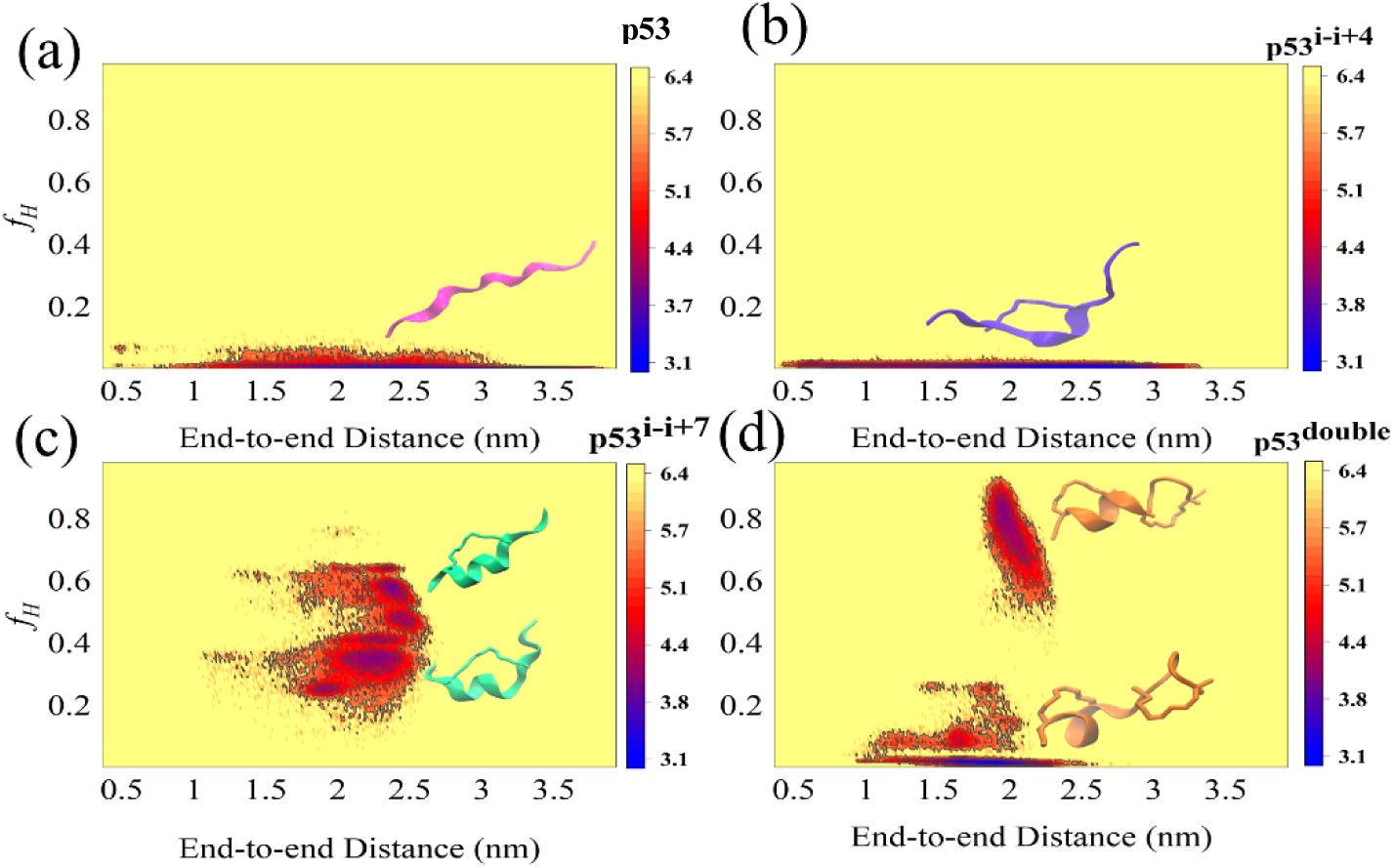
Free energy surface of end-to-end distance and the fraction of helicities of (a) wild type p53, (b) p53^i-i+4^, (c) p53^i-i+7^ and (d) p53^double^ in aqueous solution. The blue-to-red-to-yellow color scale represents increasing conformational free energy in kcal mol^−1^. The blue color area in the free energy plot corresponds to energy minima points which are highly populated whereas the yellow color represents conformations that are less populated state of peptides throughout the last 500ns unbiased simulation.

According to Figure 3, wild-type p53 and p53^i-i+4^ cover a wide range of end-to-end distances having a helicity value below 0.1. However, p53^i-i+4^ has populated basins having slightly greater helicity than p53 (Figure S3 (a)) because of a single i-i+4 cross-linker. Similarly, it has conformations with end-to-end distances in the range of 0.5nm - 3nm which is lower than wild-type p53. The end-to-end distance of p53^i-i+7^ varies approximately between 1.8nm to 2.8 nm showing the different populated basins at different end-to-end distance values. One of the larger populated basins of p53^i-i+7^ has an end-to-end distance between 2 nm to 2.5 nm whereas other basins are above 2.5nm. However, there is a small energy basin is also seen around 1.8nm. Likewise, going from p53^i-i+7^ to p53^double^ we can see the end-to-end distance covers a shorter range starting from 1nm to 2.5nm corresponding to two populated energy basins, out of which the helicity of the highly populated area is around 0.5 and for another one, it is between 0.6 to 0.8. This populated end-to-end distance for all systems is also reflected in the end-to-end distance probability plot in Figure S4.

Hence it is clear from Figure 3 that, on average the end-to-end distance of p53^i-i+7^ is greater than p53^double^, but the alpha helicity is maintained for both the peptides. Even though p53^i-i+7^ has larger end-to-end distance conformations, it is capable of maintaining high helical structures whereas p53^double^ has conformations with helicity values cantered at 0.1 and 0.7. These two types of conformations are quite distinct and conformations with helicity values intermediate between 0.1 and 0.7 barely exist. This can be seen in Figure S3(a), where two distinct peaks are obtained for p53^double^.

Hence, the length and position of the cross-linkers have a great role in deciding peptide structure. On comparing the free energy surfaces of p53^i-i+7^ and p53^double^, it is clear that end-to-end distance and helicity are not always correlated with each other. A peptide having a longer end-to-end distance can have a high helicity value. This is due to the unfolding of terminal residues of a peptide and the range of the residues covered by the cross-linker. Similarly, a peptide with a shorter end-to-end distance loses its helicity due to the breaking of the intrapeptide hydrogen bonds. This formation and breaking of intrapeptide bonds can be better explained from hydrogen bond analysis of peptides which is discussed in later section 3.4. In addition, this variation of helicity has an impact on per-residue helictites which is discussed in detail in section 3.3.

#### 3.3 Residue-wise helical occupancy of peptides in aqueous solution

In Figure 4(a), there is a pronounced difference in secondary structures for all residues of various peptides and this is calculated over the last 500ns by the STRIDE^66^ from VMD. Initially, each peptide has an alpha-helical nature, but the unfolding of these peptides occurs at different times. In the case of wild type p53, it maintains its secondary structure up to 300ns whereas for p53^i-i+4^, it is 420ns. Both p53^i-i+7^ and p53^double^ have α-helices for a longer time compared to wild-type p53 and p53^i-i+4^ but p53^double^ loses its secondary structure after 700ns by all of its residues. For p53^double^ residue ranges from Thr2 to Leu9 only show helical behavior, though two staples are attached at Thr2-Leu6 and Lys8-Glu12. This implies that p53^double^ has one half more stable than the other. On the other hand, p53^i-i+7^ residues ranging from Phe3 to Glu12 maintain their secondary structure since the i-i+7 cross-linker spans from Ser4 to Pro11 positions. The terminal residues have negligible helicity from the start of the simulations. Also, p53^i-i+7^ loses its alpha-helical secondary structure around 250-400ns of the simulation but then regains it and maintains it throughout the rest of the simulation time, whereas p53^double^ becomes unfolded after 700ns and remains coiled.

**Figure 4:**
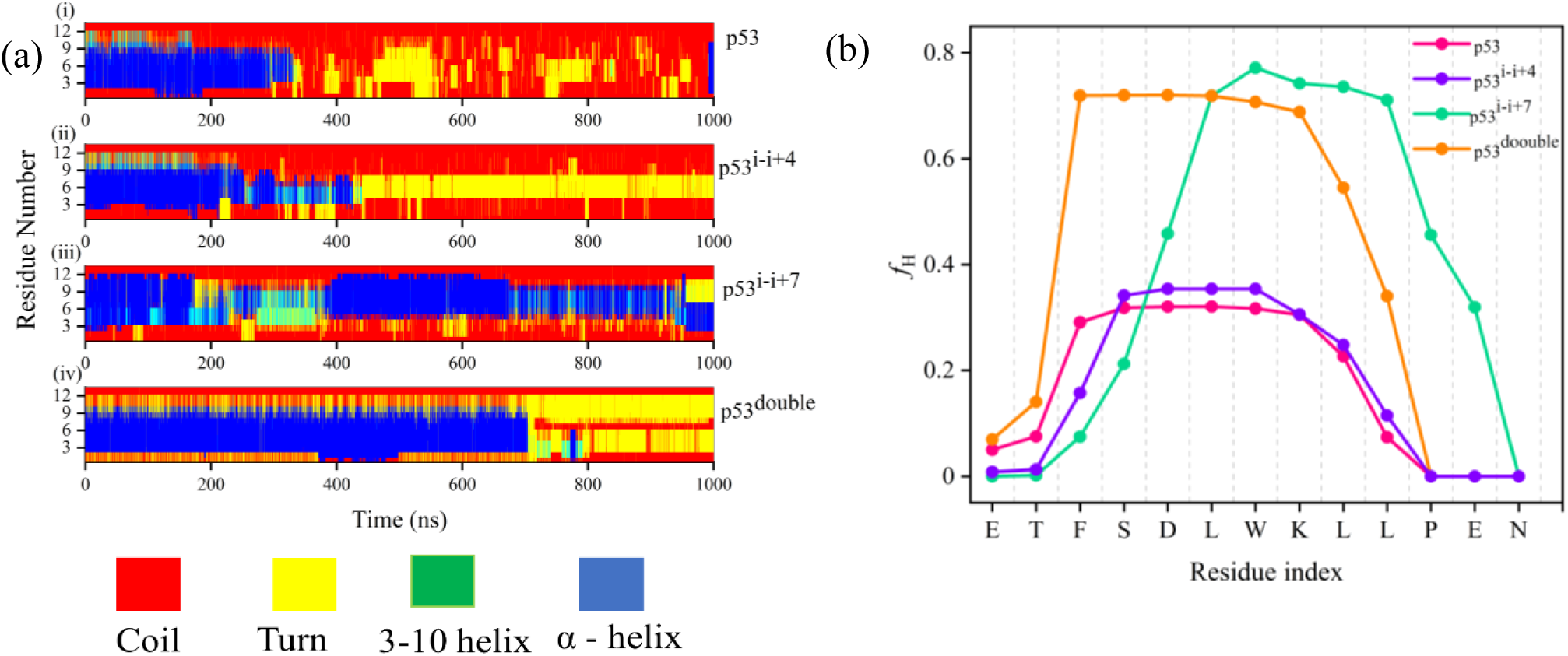
(a) Time evolution of the secondary structures of (i) p53, (ii) p53^i-i+4^, (iii) p53^i-i+7^, and (iv) p53^double^ along the 1microsecond of the trajectory. The color code for secondary structures are labeled below the graph which are as, blue → α-helix, yellow → turn, red → coil. (b) Per residue helical fractions of p53 (magenta), p53^i-i+4^ (violet), p53^i-i+7^ (green), and p53^double^ (orange) are plotted together. The four graphs depict per residue helical fraction for the last 500ns simulations.

To further quantify, we have computed the residue-wise helical fraction of all peptides in the free state in Figure 4(b). Here we found that, for each stapled peptide, those residues that are covered by the cross-linkers have high values of helicities compared to other residues. p53^i-i+7^ residues have a greater degree of helicity. For p53^i-i+7^ residues ranging from Leu6 to Leu10 have high values of *f*_H_ between 0.7 to 0.8. This is further supported by the calculation of intrapeptide H-bonds discussed in section 3.4. p53^double^ follows a similar trend of helicity as p53^i-i+7^ for different ranges of residues. The first i-i+4 cross-linker causes the higher helicity (∼0.7) values for Phe3 to Trp7 of p53^double^. Despite having a cross-linker attached at Lys8 - Glu12, the peptide loses helicity in this case, because a kink is formed between the two short length i-i+4 cross-linkers. This unusual behavior is in well agreement with the populated conformations sampled by p53^double^ in the end-to-end distance and helicity free energy surface shown in Figure 3. This unique behavior of p53^double^ can also be explained by the formation of an intrapeptide hydrogen bond between residues i and i+3 or i and i+4 residues. We discussed this in the next section.

On the other hand, p53^i-i+4^ and wild-type p53 behave similarly and have lower values of *f*_H_ than p53^i-i+7^ and p53^double^. Due to its i-i+4 cross-linker being linked in position 4 to 8, p53^i–i+4^ exhibits a slight rise in *f*_H_ from Asp5 to Lys8. This residue level helicity gives us information about the overall efficiency of the cross-linker and the importance of its position of attachment to peptide for maintaining structural stability.

#### 3.4 Intra peptide hydrogen bond between residues

The globular structure of a protein is stabilized by various factors and one of these is the hydrogen bond network. The conformational stability and helical structure of peptides are also supported by the intrapeptide hydrogen bonds particularly i-i+4 backbone hydrogen bonds (alpha-helical hydrogen bonds) (Figure 5(a)). These bonds are formed by the backbone carbonyl (C=O) group of residue i and a backbone amine group (N=H) of (i+4)^th^ or (i+3)^th^.^67^

**Figure 5:**
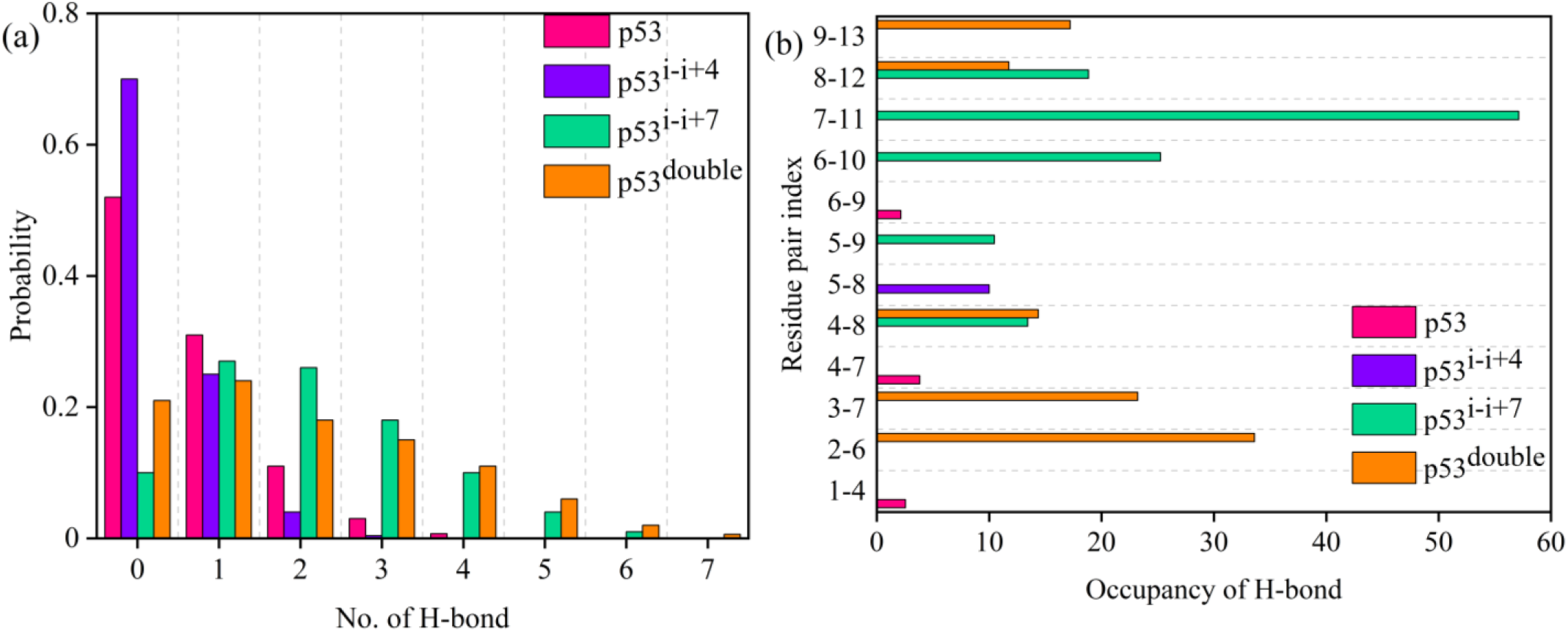
(a) Probability of intrapeptide hydrogen bonds for peptides p53, p53^i-i+4^, p53^i-i+7^, p53^double^ and (b) Percentage of occupancy of hydrogen bonds between i-i+3, i-i+4 residue pairs showing i-i+4 residue pair hydrogen bonds are highly populated for stapled peptides. These above calculations are done over the last 500ns of unbiased simulation of free peptides.

From Figure 5(b), it is clear that hydrogen bonds are highly populated at the residues where the cross-linker is attached and its nearby residues. For instance, in the case of the p53^double^, the most hydrogen bonds are formed between the stapled residues Thr2 and Leu6 and Phe3 and Trp7. Owing to the above reason, these residues also possess a high value of *f*_H_ described earlier in section 3.3. Moreover, these hydrogen bonds have an occupancy of around 35% and 25 % respectively in the last 500ns simulation. In addition to this, there is also a hydrogen bond found between 4Ser and Lys8 of p53^double^ which is responsible for maintaining the helicity at Trp7 and Lys8 of p53^double^ shown in Figure 4. In addition to this, there are two more H-bonds found between the pair of residues Lys8-Glu12 and Leu9 -Asn13 due to the spanning of the second cross-linker from Lys8 to Glu12. However, these two H-bonds are very less stable having occupancies below 20% so this part of the peptide gradually loses its helicity as shown in the residue-wise helicity plot (Figure 4 (b)). Also, due to the absence of H-bonds for residues lying in between the two i-i+4 cross-linkers, there is a kink obtained for p53^double^ and is reflected in the smaller end-to-end distance of p53^double^ and low value of helicity fraction (*f*_H_) (Figure 3).

On the other hand, for p53^i-i+7^, several hydrogen bonds are found with greater lifetimes due to the span of its long i-i+7 cross-linker between the Ser4 and Pro11 stapled positions. Among all these H-bonds, the highly populated H-bond is found in between Trp7 and Pro11 with an occupancy of around 58 – 60 %. Because there are more frequently occurring adjacent H-bonds between the i^th^ and i+4^th^ residues of p53^i-i+7^, it preserves the overall secondary structure with all of its residues having a larger fraction of helicities. Contrary to p53^i-i+7^, p53^i-i+4^ behaves like wild-type p53 despite having one shorter length staple that spans over fewer residues. As a result, only one i-i+3 H-bond is formed between Asp5 and Lys8. Here, wild-type p53 and p53^i-i+4^ exhibit a very low occupancy of i-i+3 H-bonds, thereby losing their secondary structure which results in a higher conformational entropy value described in section 3.9.

### Replica exchange molecular dynamics simulation

To explore the conformational distribution of the stapled peptides, one of the enhanced methods, T-REMD^35^ has been used and the structural behaviors of peptides have been investigated.

#### 3.5 Convergence of Replica exchange molecular dynamics simulation

In the T-REMD^35^, the peptide is simulated in several copies simultaneously, each at a different temperature periodically swapping their configurations with nearby replicas. The convergence of the aforementioned method can be estimated from the potential energy overlapping of two nearby windows. Here, we have calculated the potential energy for the first 10 replicas for each peptide (Figure S5) which shows a significant overlap of the potential energy between adjacent windows. In addition to this, a fair exchange probability of conformations is found (between 0.35 and 0.40) whereas the desired exchange probability is set as 0.25. This implies a single replica conformations explore possible temperature space. The exchange probabilities for the first 10 replicas for each studied peptide are given in supporting information (Table ST1).

#### 3.6 Free Energy Surface along Rg and RMSD from T-REMD

T-REMD is the potential enhanced sampling method to access the conformational space beyond the stipulated timescale.^35^ The replicas simulated at different temperatures access those conformations and provide fair inference about the free energy landscape. In Figure 6, the free energy surface emphasizes that the addition of a cross-linker along the sequence substantially restricts conformational exploration mapped from the lowest temperature index replica. In Figure 6, the wild-type p53 and p53^i-i+4^ show comparable behavior at greater variations in Rg and RMSD. However, with the introduction of a small i-i+4 cross-linker, the populated basins are shifted towards low Rg values, though RMSD variation still occurs at higher values for both of the peptides. The conformational space restriction is significant in p53^i-i+7^ and p53^double^ (Figure 6 (c) and Figure 6 (d)) compared to wild-type p53 and p53^i-i+4^. The variation in Rg for p53^i-i+7^ is less than the p53^double^ whereas for p53^double^, variation in RMSD is limited. Nevertheless, the populated basins are obtained at higher RMSD in the case of p53^double^. Here, p53^double^ samples a large number of conformations at a high RMSD range compared to its unbiased simulation (Figure 2(j)). However, the relative trend in the free energy landscape (Figure 6) is concurrent with the unbiased simulation results in Figure 2 (a), (d), (g), and (j). The superimposed plots of these free energy surfaces are shown in supporting information (Figure S6). This infers a larger exploration of the free energy surface for T-REMD simulation over the unbiased simulation.

**Figure 6:**
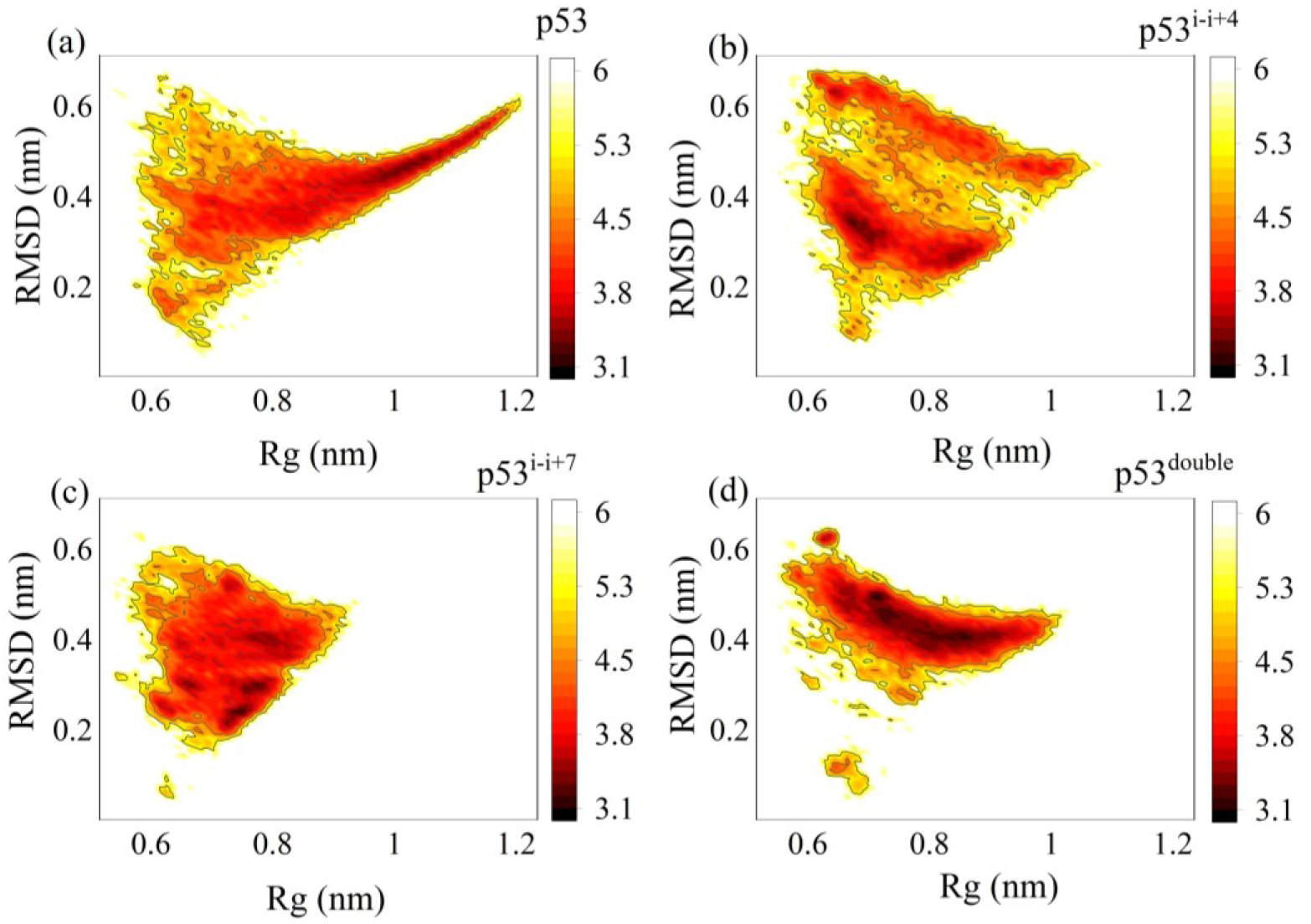
Free energy surface along Rg and RMSD obtained from T-REMD simulations for systems (a) p53, (b) p53^i-i+4^, (c) p53^i-i+7^, and (d) p53^double^ are represented with energy color bars. Free energy values are in kcal mol^−1^.

#### 3.7 Exploration of peptide conformation via Clustering

T-REMD involves generating trajectories of multiple conformations having similar or different structural properties. Among the various structural properties, Root mean square deviation (RMSD) is one of the important measures that can differentiate conformations from each other. To examine various peptide conformational states and classify related conformations, RMSD-based clustering can be employed. All these clusters are categorized by the GROMOS algorithm^68^ and the RMSD cutoff has been set to 0.2 nm. According to this algorithm, the number of neighbors of each structure has been assigned to a particular cluster using the specified RMSD cutoff value. This structure and its neighbors are then eliminated from the pool of conformations. Further distinct clusters are assigned with respect to the RMSD cutoff until all the rest conformations have been assigned to a cluster. This clustering scheme helps us to identify key conformations.

Here, in our work from Figure 7(a), we depict that, the number of clusters varies from one system to another. This establishes that the addition of cross-linker is found to be effective in maintaining the conformational rigidity of the peptide. Among the stapled peptides, p53^i-i+4^ shows a larger number of clusters. The p53^i-i+7^ has comparable numbers of clusters with p53^double^. However, only the number of clusters can’t give detailed information about the conformational exploration of a peptide. Therefore, to look into the detailed picture, the populations of the first five clusters are analyzed. Here clusters are sorted in increasing order of their RMSD values. In Figure 7(b), the p53^i-i+7^ exhibits with highest number of populations in the first cluster implying the least RMSD deviation in overall conformational space while p53^double^ follows a comparative trend with slightly more distributed conformation over the top five clusters. This implies p53^double^ shows less conformational variation along RMSD which is reflected in its free energy surface (Figure 6(d)).

**Figure 7:**
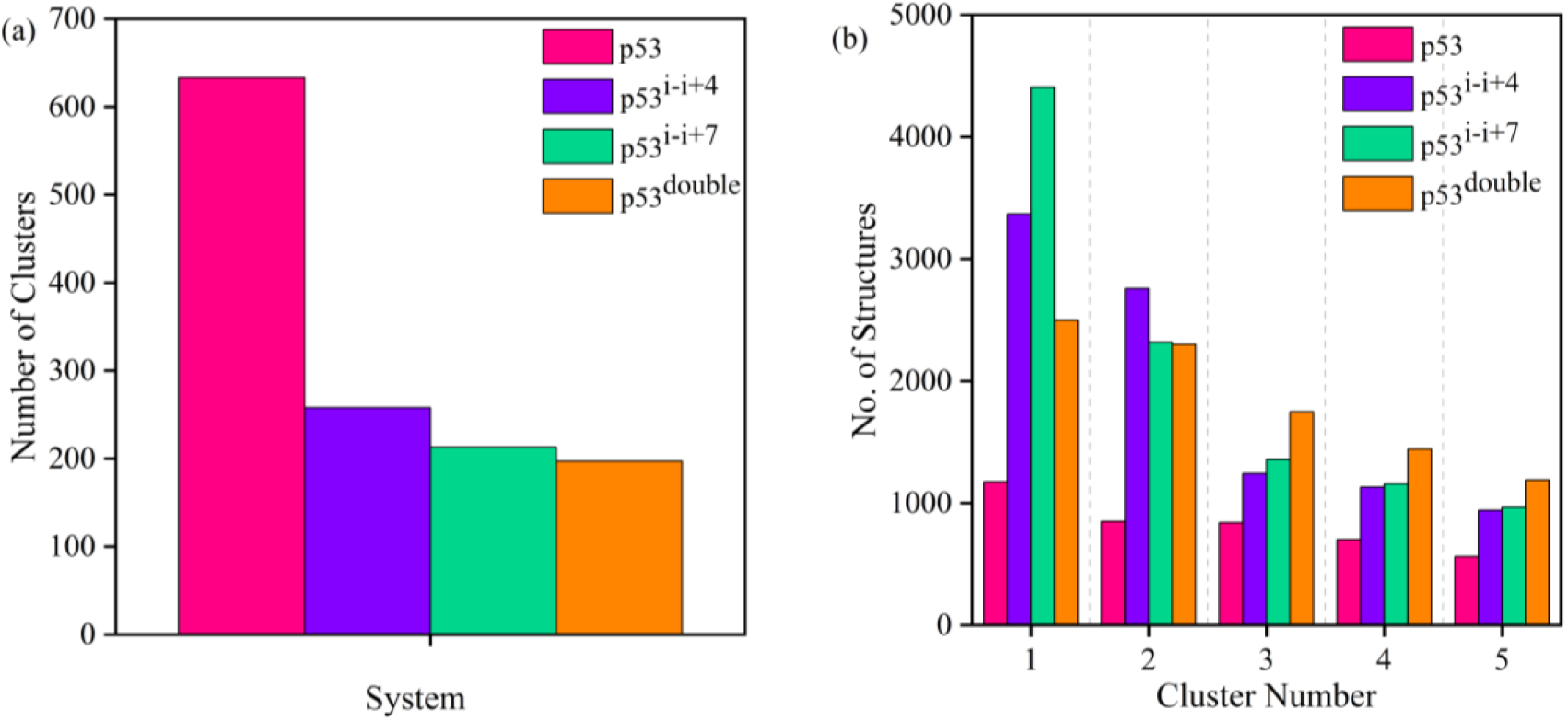
(a) Number of clusters computed for p53, p53^i-i+4^, p53^i-i+7^, and p53^double^ from T-REMD simulation of 50ns in each replica (b) Population of top 5 clusters for each system. The color of the system is represented in the Figure.

#### 3.8 Helicity melting curves

To gain a further understanding of the conformational behavior of free peptides, these are investigated in high-temperature space through a temperature-replica exchange simulation. Here, the persistence of the peptide’s helical structure is investigated as a function of temperature. From this helicity melting curve (Figure 8), it is clear that with the increase in temperature, the helicity of all peptides is affected. To begin with, at 310K, all of these systems have comparative helical content. All the stapled peptides have slightly higher helicity than wild-type p53 at physiological temperature.

**Figure 8:**
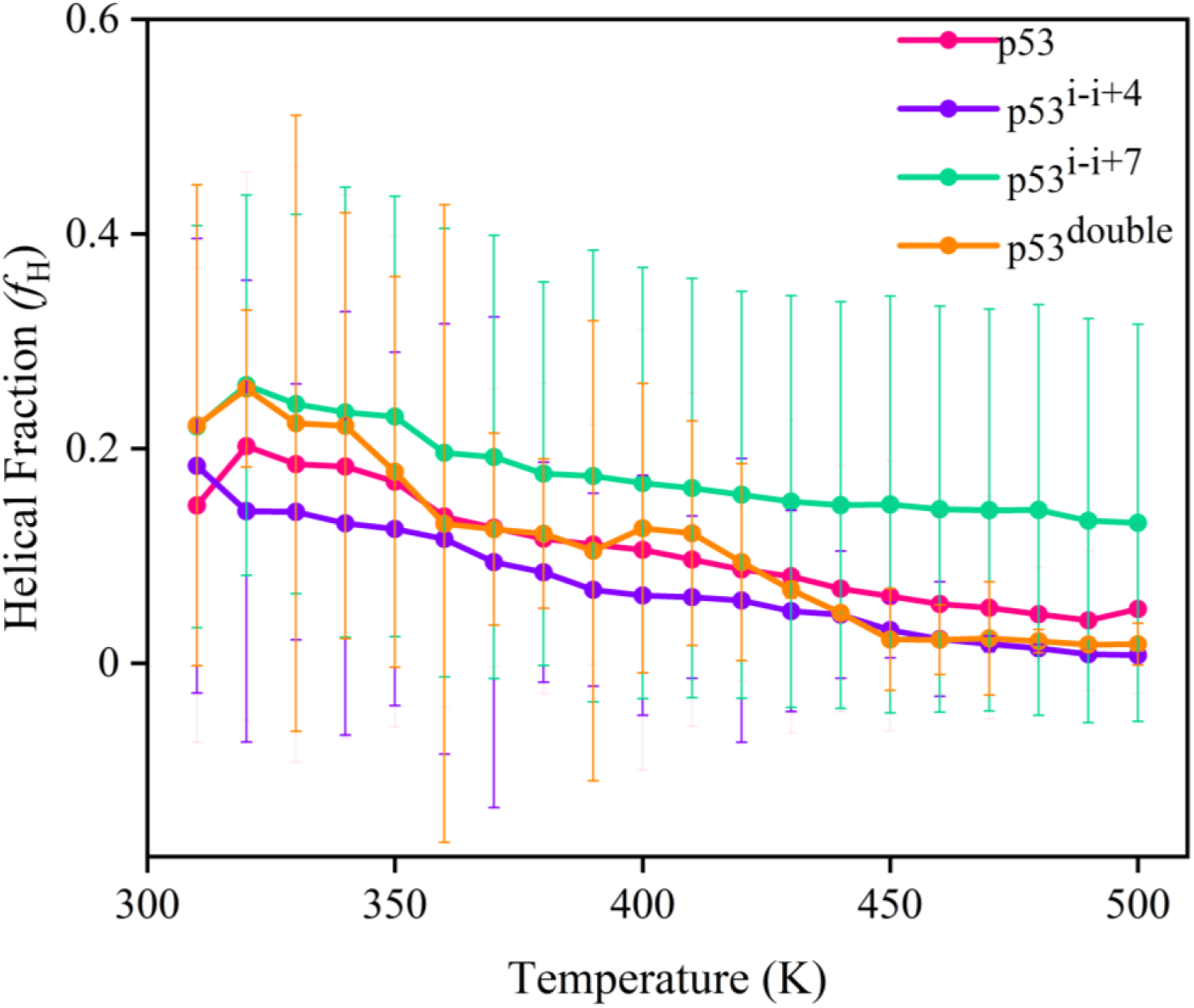
Helicity melting curves for p53, p53^i-i+4^, p53^i-i+7^, and p53^double^ from T-REMD simulation are shown with their respective error bars. The color code for each system is represented in figure legends.

In Figure 8, the p53^i-i+7^ maintains the helical content intact over the large temperature range while other peptides tend to unfold with increasing temperature. The helical fraction remains near 0.2 for the p53^i-i+7^. In the case p53^double^, the initial low temperature replicas have significant helical content, but in the high temperature, it unfolds in parallel with p53^i-i+4^. In a nutshell, p53^i-i+7^ persists at comparatively greater helical fractions even at high temperatures while other peptides show decay in helical fractions. This superiority of p53^i-i+7^ has also been supported from the helicity probability distribution from unbiased simulation (Figure S3(a)). It has been also reflected in end-to-end-distance and helical fraction free energy surface generated from T-REMD simulation (Figure S7). The helicity melting curve is plotted with the average helical fraction from the respective temperature replicas. The error bars showing high standard deviations arise from the swapping of conformations from the adjacent temperature of T-REMD simulations.

Overall, in unbiased molecular dynamics and temperature-replica exchange molecular dynamics studies, stapled p53 peptides sample less flexible conformations compared to wild p53 peptides in the free state. This conformational flexibility is highly dependent on the choice of cross-linkers such as its chemical nature, length, and position of attachment to the peptide. In our study, we observed that while p53^double^ has more staples, it cannot sustain its secondary structure due to the spanning of shorter cross-linkers over the sequence and their point of attachment. On the other hand, single stapled p53^i-i+7^ maintains its helicity. This conformational rigidness will affect the entropy of peptides in the free state and the bound state with MDM2. Hence, the investigation of the thermodynamics of p53 and its stapled versions with MDM2 be extremely useful in elucidating the driving force behind the peptide-MDM2 binding.

### Binding of p53 peptides with MDM2

#### 3.9 Enthalpy and Entropy of Stapled Peptides

In this section, we reported the calculated values of binding free energies, considering both enthalpy and entropy. We have calculated the enthalpy of binding between p53 and MDM2 using the MMGBSA protocol implemented in the GROMACS package.^62^ The details of this method have been discussed in section 2.4. This method has been earlier used in describing the binding free energy between protein and peptide successfully even in the case of stapled peptides with MDM2.^34,63^ On the other hand, the entropy is calculated using the quasi-harmonic approximation.^69^ Both enthalpy and entropy have been calculated on the conformations from the last 500ns of simulations. Here, the multiple simulations of the MDM2+p53, MDM2+ p53^i-i+4^, MDM2+p53^i-i+7^, and MDM2+p53^double^ systems are carried out where MDM2 is the receptor and p53 is the peptide ligand. Since we are using a multi-trajectory Protocol (MTP)^61^ for the binding free energy computation, three independent trajectories for MDM2 and p53 peptide are also taken. The values for individual simulations are given in supporting information tables (ST2 to ST5).

In Table 3, it is apparent that all of those 3 stapled peptides have favorable binding free energy than wild-type p53 with MDM2. When a cross-linker is added to a peptide, the enthalpy of the interaction between the surface of the binding pocket and peptide changes. In addition to this, the entropic penalty to complex formation also varies since the peptide undergoes conformational changes from its free state to bound state. Our calculation shows that p53^double^ has higher binding affinity with MDM2 than other stapled peptides. However, the contribution of enthalpy and entropy to binding free energy differs from one stapled peptide to another. For p53^i-i+7^, both the enthalpy and entropy change are less compared to other peptides. In contrast, for p53^i-i+4^ and p53^double^, the enthalpy change is larger due to their interactions at the binding surface of MDM2 resulting in more conformational changes of those peptides in the bound state than the free state giving a larger entropy change. For wild-type p53, the binding free energy determined by MMGBSA is in good agreement with the range of experimentally observed binding free energy values, which are in between −6.8 and −9.1 kcal mol^−1^.^15,37,70^ In addition to that, a single-aliphatic i-i+7 stapled p53 peptide investigated experimentally has shown to have a higher binding affinity with MDM2 than wild-type p53 peptide.^30^ Several experimental investigations have demonstrated the usefulness of double-stapled peptides in preventing protein-peptide interactions by having higher cell permeability, proteolytic stability, etc. For instance, a peptide containing double i-i+4 hydrocarbon cross-linkers has been demonstrated to be proteolytically stable and a good HIV-1 fusion inhibitor.^71^ Recently a p53 derived double-stapled peptide has been found effective in maintaining helicity, cell permeability, and proteolytic stability and can act as a potent dual inhibitor for p53-MDM2/MDMX interaction.^72^

Since binding free energy consists of enthalpy and entropy, there is always a tug of war between these two components. Hence, a component wise decomposition of overall binding free energy is important. The difference in enthalpy between the free state and bound state of a complex arises due to both gas phase energy and solvent phase energy. In general, the gas phase energy is mostly influenced by electrostatic and van der Waals energy whereas solvent phase energy is contributed by the polar solvation and non-polar solvation. From the binding free energy values (Table 3), there is a significant difference between wild-type p53 and stapled peptides, with the electrostatic component changing by around 40 kcal mol^−1^ and the solvation energy changing by about 50 kcal mol^−1^. The single stapled peptide p53^i-i+7^ has a lower enthalpy value than wild-type p53 despite having an i-i+7 cross-linker due to the loss of electrostatic and van der Waals interaction whereas both p53^i-i+4^ and p53^double^ shows an enthalpic gain binding free energy upon cross-linking. Although key components like electrostatic, van der Waals, polar solvation energy, etc. have comparable values between p53^i-i+7^ and p53^double^, there is also a significant difference between their enthalpy value, which is around 30 kcal mol^−1^. This variation in enthalpy arises from different 1-4 electrostatic interactions and some of the internal energy components, including dihedral terms and angel terms. This is attributed to the different orientations of the stapled peptide residues on the MDM2 surface and their altered conformations in the bound state relative to the free state This can be better understood by looking into the residue-wise decomposition of enthalpy (section 3.10). Similar to p53^double^, p53 with a single i-i+4 staple shows larger enthalpy change as a result of orientation changes of residues in the bound state than in the free state.

The entropy loss shows that each peptide prefers a rigid conformation in its bound state over a free state (Figure S8). However, each of the stapled peptides has less entropy than the wild-type in the free state. This type of change in conformational entropy upon backbone rigidification has been seen in cyclic peptides from linear peptides.^73^ From Table 3, p53^i-i+7^ shows less entropy change compared to p53^i-i+4^ and p53^double^ upon complex formation with MDM2. This less entropy loss for p53^i-i+7^ can be explained by the persistence of alpha-helical structures in the free state. Further, the increase in entropy loss for p53^i-i+4^ and p53^double^ can be explained by the helicity overlapping plot of these peptides (Figure S8). From Figure S8, there is a larger helicity difference is seen for p53^double^ between its complex state (grey color lines) and free state (orange color lines) in the last 300 ns of simulation. This lower efficiency of free p53^double^ to form an alpha helical structure is due to its short length cross-linkers and their positions of attachments to the peptide. All of these conformational dynamics studies, we have discussed earlier. (sections 3.1-3.9). Furthermore, the rigid conformation of the p53^double^ in the bound state is an outcome of its orientation and interaction of residues with MDM2. Thus, there is a considerable conformational change in the bound state than the free state resulting in a larger entropy penalty. A similar type of behavior is seen in the p53 peptide having a single i-i+4 aliphatic cross-linker (p53^i-i+4^). A visual representation of these rigid conformations of the p53 peptide in the bound state is added in the supporting information (Figure S8). A residue-wise decomposition of enthalpy will give a clearer picture of interaction with MDM2, discussed in section 3.10.

#### 3.10 Decomposition of Enthalpy

The residue-wise decomposition of enthalpy gives information about the feature of cross-linkers and peptide sequence and its binding with the target partner. To further understand the above binding free energy values in detail and the role of each residue in binding, different components of binding energy have been decomposed for each residue in the following section.

In Figure 9, it is observed that every residue of p53 has varied contributions to the binding with MDM2. Overall, p53^i-i+7^ has less enthalpy contribution to its binding with MDM2 whereas the other two stapled peptides show high enthalpy contribution (Table 3). This enthalpy contribution comes from different energy components such as electrostatic, van der Waals interaction, and polar and non-polar solvation energy as discussed in the binding free energy calculation section (section 2.4).

**Figure 9:**
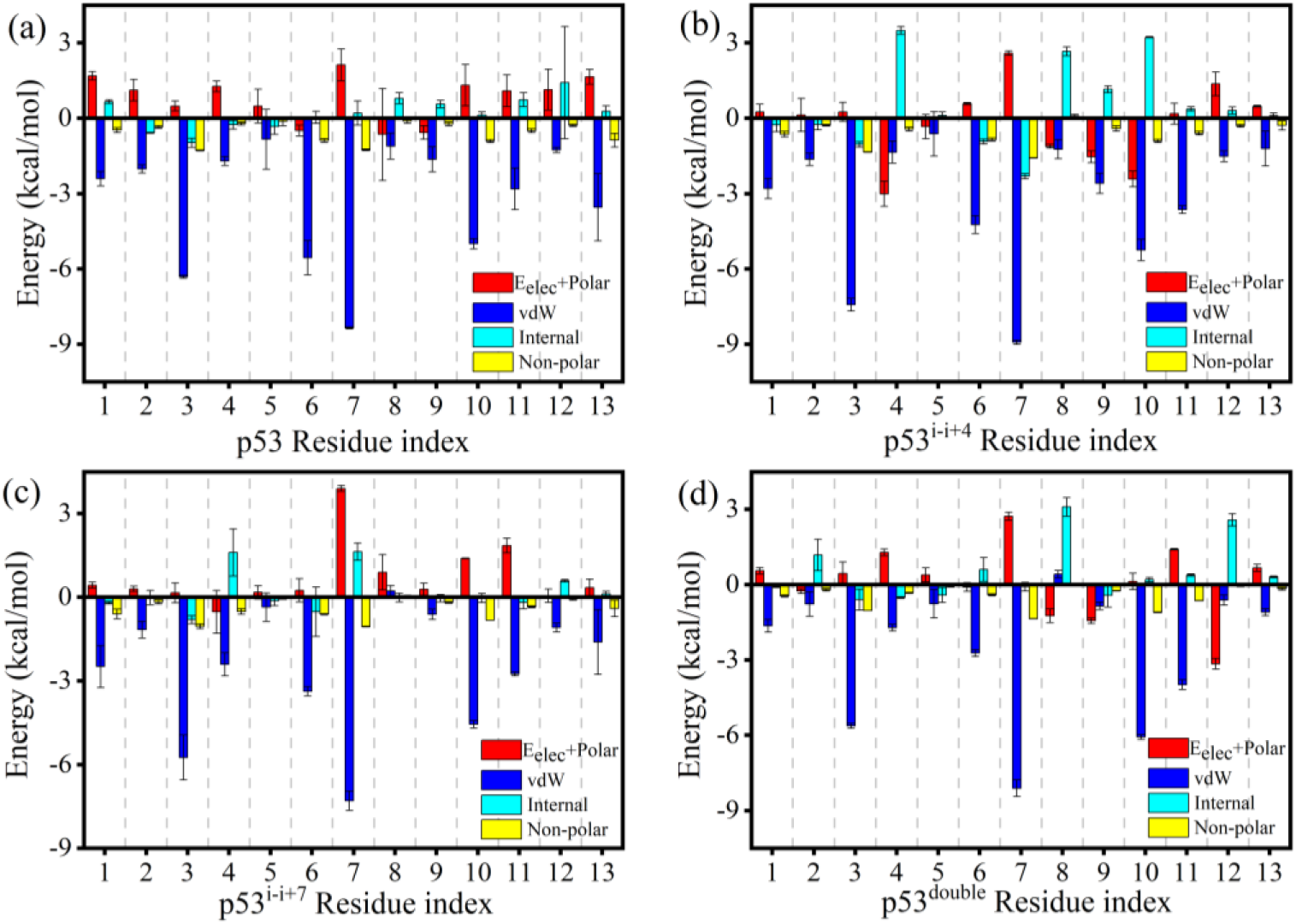
Decomposition of enthalpy to different energy component contributions such as electrostatic energy, polar solvation energy, van der Waals interaction, internal energy, and non-polar solvation energy for systems (a) MDM2 + p53, (b) MDM2 + p53^i-i+4^, (c) MDM2 + p53^i-i+7^ and (d) MDM2 + p53^double^ are represented. The respective color codes of energy components are the same and are mentioned in their corresponding Figure legends. This decomposition of residues is averaged over the last 500ns of 3 independent simulations and respective error bars are shown in the above Figure.

In wild-type p53 (Figure 9(a)), the enthalpic contribution for binding of wild-type p53 with MDM2 mainly arises from van der Waals interaction. On the contrary, it shows a positive value of the sum of electrostatic and polar solvation energy contributions for most of the residues. So, here this unfavorable electrostatic and polar solvation energy is compensated largely by van der Waals interaction, and by a small amount of non-polar solvation energy and internal energy of the peptide, as a result, the enthalpic gain is observed.

From section 3.9, it can be shown that adding a cross-linker to peptide results in a substantial decrease in the binding free energy of p53-MDM2. Further, the component-wise energy contributions vary depending on the type of cross-linker attached to the sequence. When p53 is attached with a single i-i+4 aliphatic cross-linker, it exhibits change in enthalpy than wild-type (Table 3). Then per-residue decomposition of this enthalpy change shows both the polar and non-polar interactions are responsible (Figure 9(b)). While wild-type p53 shows the unfavorable positive value of electrostatic and polar solvation energy contribution for its residues, for p53^i-i+4^, some of the residues have favorable polar interaction energies. Moreover, p53^i-i+4^ has some significant differences in internal energy components explaining the peptide residues undergo structural arrangements in bound state for favorable binding. From Table 3, it is clear that although p53^double^ has effective favorable binding free energy with MDM2 than p53^i-i+4^, the energy contributions to enthalpy change are almost comparable (Figure 9(d)) with p53^i-i+4^. Similar to p53^i-i+4^, p53^double^ residues contribute to van der Waals interaction and non-polar interaction. Also, some of its residues contribute to electrostatic energy. In addition to this, its residues show a large change in internal energy from the free state to the bound state. This arises due to their structural arrangement and orientation at the MDM2 binding surface which is also responsible for larger entropic loss during complex formation. Hence, p53^double^ shows a larger enthalpy change during binding with MDM2.

On the other hand, p53 with an i-i+7 aliphatic cross-linker exhibits lower enthalpy than the wild-type. This is explained by the changes in the individual energy components. The van der Waals interaction of p53^i-i+7^ is weaker than that of the wild-type p53. It has also a decrease in electrostatic and polar solvation energy due to the comparatively smaller number of electrostatic contacts. However, p53^i-i+7^ shows a favorable binding free energy with MDM2 than wild-type p53. This is explained by the ability of p53^i-i+7^ to form rigid structures in the free state and thereby modulate the formation of the complex with MDM2 by reducing the entropic cost.

A comparison of the orientations of the wild-type p53 and the single and double stapled p53 peptides in the binding surface can be well understood from the visualization of the simulations of MDM2+p53 (Movie S1), MDM2+p53^i-i+7^ (Movie S2) and MDM2+p53^double^ (Movie S3). The difference in orientation of the p53^i-i+7^ and p53^double^ residues with MDM2 results in distinct populations of hydrogen bonds (Figure S9). Therefore, the nature, length, and position of the staple on the peptide determine the binding free energy of these peptide inhibitors with their target partner.

## 4. Conclusions

In the present study, we have used extensive all-atom molecular dynamics simulations to study the structural distributions of several cross-linked peptide analogs belonging to the transactivation domain of p53. Though a variety of cross-linkers are usually used to design such analogs, hydrocarbon cross-linkers have been proven successful for their helix inducing properties. In this article, we provided a thorough description of the impact of the hydrocarbon cross-linker on the conformational dynamics of peptides and the thermodynamics of binding by altering the length and position of the cross-linker’s attachment to the peptide. We began our study by considering designing a stapled p53 peptide with an i-i+4 aliphatic cross-linker and then used computer simulations to study the impact of the cross-linker on the conformational dynamics of the peptide. Due to its shorter length, the cross-linker is unable to cover all the residues across the peptide sequence. In order to make a comparison, we modeled two stapled peptides, one includes a single i-i+7 cross-linker and the other has two i-i+4 cross-linkers. Cross-linking, in general, restricts the conformational dynamics of the peptides. The conformational landscape has been calculated by taking into account two structural parameters viz. Rg and RMSD. For cross-linked peptides, the restricted motion of the backbone atoms results in smaller changes in RMSD values. The restricted motion is also responsible for maintaining the overall helicity of peptides in the free state. To test the effectiveness of the stapling agents, two perturbing conditions are imposed i.e. thermal and chemical. In one case the temperature is raised to 330K and in the other case 8M urea is added. It is found that, indeed stapled peptides can limit conformational sampling even at perturbing conditions. The overall propensity to form alpha-helical structures seems to vary between p53 peptide analogs and it is dependent on the chemical nature of the staples and their position of attachment to the peptide. From the comparison of the stapled peptides in denaturing conditions, p53 with a single i-i+7 cross-linker is found to be more effective than the rest of the stapled peptides and p53^i-i+4^ behaves similarly to the wild-type p53. This efficiency of p53^i-i+7^ is also studied by computing the free energy surface along end-to-end distance and helicity. The free energy surface infers that though it has a larger end-to-end distance than p53^double^, due to the unfolding of terminal residues, it has a high helical fraction throughout the simulation. The populated conformations in p53^i-i+7^ have a continuous increase in helicity while p53^double^ shows distinctly populated regions with two discrete helicities. Owing to the absence of a cross-linker on the middle residue of the p53^double^, it adopts a kink. This kink-type structure affects the p53 ^double^ secondary structure and the per residue helicity. The secondary structure analysis is also supported by intrapeptide hydrogen bonds, particularly alpha-helical hydrogen bond (i-i+4) populations. To accelerate the conformational distribution, we have performed an enhanced sampling method, Temperature replica exchange molecular dynamics (T-REMD) for the temperature range of 303 to 500K. The conformational flexibility and secondary structure of wild-type p53 and its designed peptides have been studied using these T-REMD simulations, and it has been observed that p53 with a single i-i+7 cross-linker is more efficient in maintaining its rigidity and helicity than the other systems. Further, to investigate the binding efficiency of these aforementioned stapled peptides with MDM2, we have taken into account these peptides in complex with MDM2. The introduction of cross-linker to peptides favors better binding with MDM2 over the wild-type peptide. This is quantified by measuring two important thermodynamic quantities, enthalpy and entropy. Enthalpy comparisons between two single stapled peptides i.e. p53^i-i+4^ and p53^i-i+7^ show that p53^i-i+4^ went through an enthalpy gain, whereas p53^i-i+7^ experienced an enthalpy loss as compared to wild-type p53 (Table 3). With MDM2, the cross-linker in p53^i-i+7^ causes the peptide to lose its electrostatic and polar solvation energy but aids in van der Waals contact due to hydrophobic interaction. On the other hand, the limited variation in the conformation of p53^i-i+7^ in both the free state and bound state leads to a decrease in entropic penalty for binding with MDM2. Conformational restrictions of p53^i-i+7^ are confirmed from both unbiased simulation and T-REMD simulations in the free state. Moreover, the binding free energy between p53^i-i+4^ and MDM2 is comparable to that of p53^i-i+7^. p53^i-i+4^ shows enthalpy gain from the free state to the bound state due to its structural arrangements and orientations of residues in the MDM2 binding surface. Since, p53^i-i+4^ exhibits more flexible conformation than p53^i-i+7^ in the free state, so it exhibits substantial entropy loss during binding with MDM2. Further, to investigate the effect of two i-i+4 stapled p53 binding with MDM2, the binding free energy of p53^double^ has been calculated. Similar to p53^i-i+4^, it also shows a large enthalpy change during the binding with MDM2. The enthalpy gain and entropy loss for stapled peptides are reflected in the binding free energy calculation with MDM2 and also in the decomposition of binding free energy. Overall, the binding free energy order is ΔG _MDM2 + p53_^double^ < ΔG _MDM2 + p53_ < ΔG _MDM2 + p53_ < ΔG _MDM2 + p53._ The effective binding free energy of double stapled peptide with a target partner is also in good agreement with experimental studies.^72^ This interplay between enthalpy and entropy change determines the overall binding efficiency. This type of enthalpy and entropy compensation has been observed in previous studies as well.^74^

Overall, our analysis suggests that the position and length of the cross-linker alter the contribution of enthalpy and entropy change to the binding free energy. This study helps in the strategic design of peptide therapeutics against MDM2.

## Supporting information

Trajectory-Videos

## Conflicts of interest

The authors declare no conflicts of interest.

## Acknowledgments

R.C. acknowledges SERB (Grant CRG/2020/000279) for financial support. A.R.C. thanks the Department of Science and Technology Inspire (DST-INSPIRE), Government of India, V.G. thanks the Council of Scientific and Industrial Research (CSIR), Government of India and thanks for Department of Biotechnology (DBT), Government of India for providing fellowship. The authors acknowledge the computational facility (HPC) provided by the Indian Institute of Technology Bombay.

## Notes

### Competing Interest Statement

The authors have declared no competing interest.

