## Supporting Information for "Exploring the effect of Hydrocarbon Cross-linkers on the Structure and Binding of Stapled p53 Peptides"

#### 1. Radial Distributions of Urea around p53

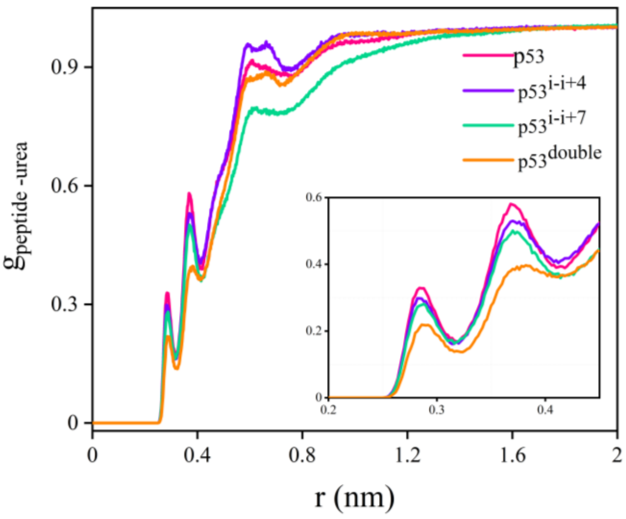


**Figure S1:** Radial Distribution function of urea around the peptide backbone for different systems p53, $p53^{i\text{-}i+4}$, $p53^{i\text{-}i+7}$, and $p53^{double}$.This RDF is calculated considering the center of mass of the urea molecule in the last 500ns of 1 μs unbiased simulation. $p53^{double}$ has less RDF compared to other systems because of less exposure of it to urea molecules.

#### 2. Residue level fluctuations

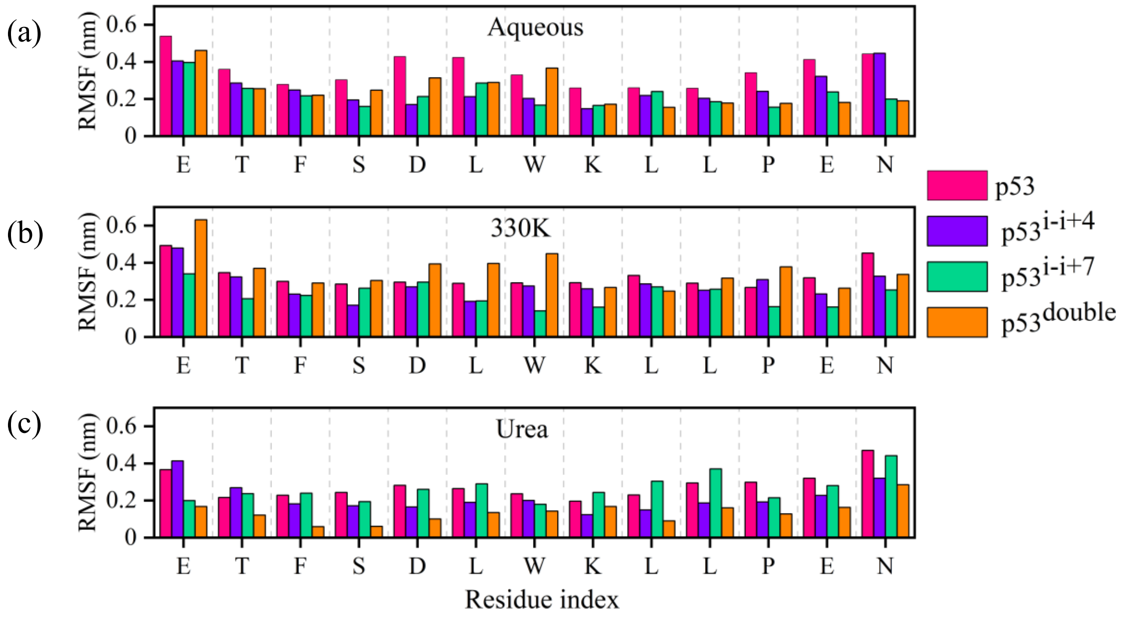


**Figure S2:** Root mean square fluctuation (RMSF) of residues of p53, $p53^{i\text{-}i+4}$, $p53^{i\text{-}i+7}$, $p53^{double}$ are presented in the bar diagram in all the 3 conditions. For system identification, the color of the histogram is maintained as the system color i.e. magenta for p53, violet for $p53^{i\text{-}i+4}$, green for $p53^{i\text{-}i+7,}$ and orange for $p53^{double}$.

Fluctuations of residues of single i-i+7 ($p53^{i\text{-}i+7}$) and double i-i+4 ($p53^{double}$) stapled peptides are much less compared to wild-type p53. This behavior is observed in both aqueous solution and denaturing conditions, high temperature (330K) (Figure S2-(b)) and chemical denaturation (8M urea) (Figure S2-(c)). Also, for a particular condition, those fluctuations of those residues are smaller which are spanned over by crosslinker. Above all this, the stapled residues show a smaller deviation compared to terminal residues in each stapled system which reflects the rigidness of residues in the presence of the stapling agent.

#### 3. Probability Distributions of Helical Fractions

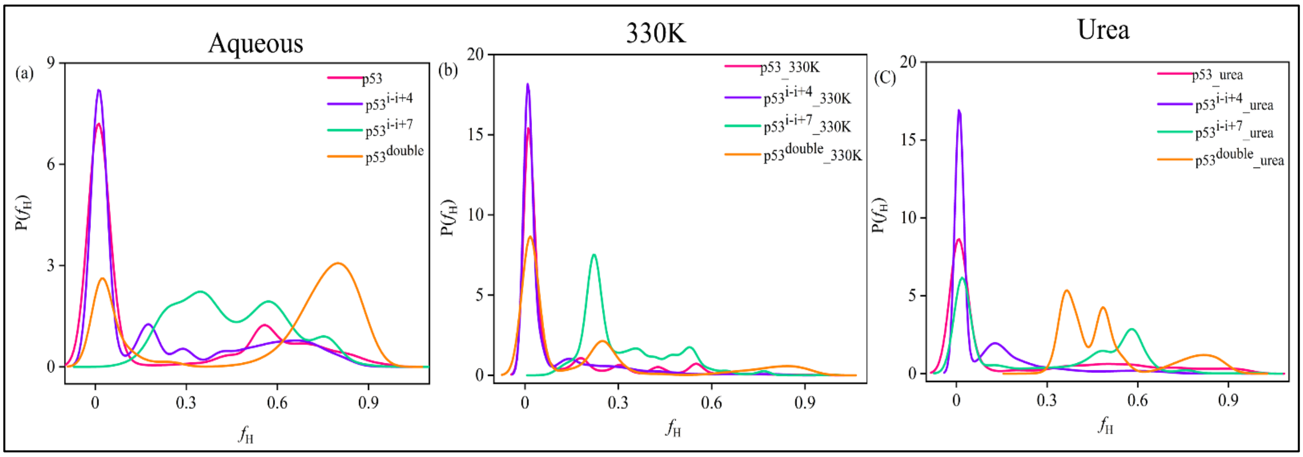


**Figure S3:** Probability distribution of the helical fractions helical fractions ($P(f_H)$) of p53, $p53^{i\text{-}i+4}$, $p53^{i\text{-}i+7,}$ and $p53^{double}$ (a) aqueous solution, (b) at high temperature 330K and (c) in 8M urea solution are represented.

#### 4. End-to-end distance probability distributions

Though here all considered peptides have equal no of residues i.e. a 13 residue peptide, but due to change in the length of the crosslinker and their attachment position on the peptide, their length varied from each other to a great extent.

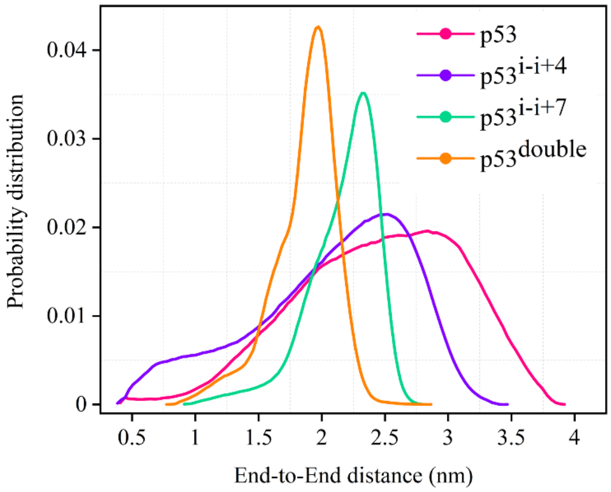


**Figure S4:** Probability distribution plots of End-to-end distance of p53, p53$^{i-i+4}$, p53$^{i-i+7,}$ and p53$^{double}$ in the last 500ns of simulation.

From this graph, it is clear that both, the wild-type and single i-i+4 p53 peptide cover a wide range of end-to-end distance that varies from 0.5nm to 4nm and 0.5 nm to 3.5 nm respectively. On the other side, p53$^{i-i+7}$ and p53$^{double}$ cover a range of 1nm to 3nm and 0.8nm to 3nm respectively. However, the probability of having an end-to-end distance between 1.5 to 2.5 nm is greater for p53$^{double}$ compared to the p53$^{i-i+7}$ peptide. So, the highest probability peak for each peptide shifted to the right side and the curve became wider when we go from p53$^{double}$ to wild-type p53 in the sequence of p53$^{double}$, p53$^{i-i+7}$, p53$^{i-i+4}$, and p53.

### 5. Potential energy overlap between adjacent replicas

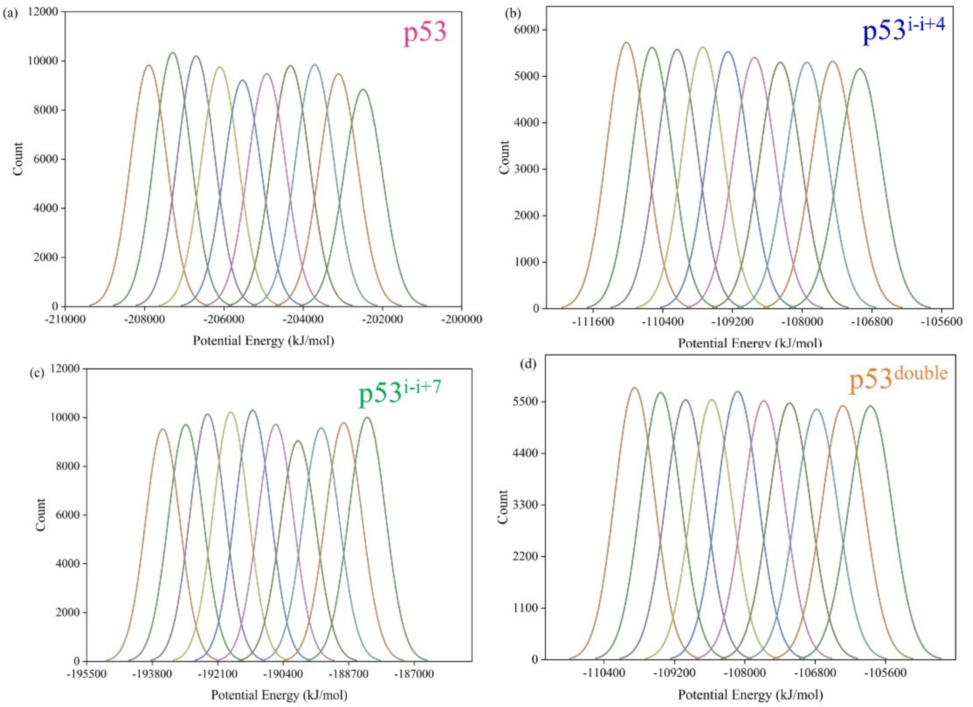


**Figure S5:** Overlapping of Potential energy probabilities for the first 10 replicas generated from Temperature replica exchange molecular dynamics (T-REMD) simulations. A significant overlapping between adjacent replica potentials shows a good exchange of conformations between the replicas implying the convergence of our T-REMD simulations.

### 6. Table ST1: Exchange probability acceptance ratio for T-REMD trajectories

| Replica ID | p53 | $p53^{i-i+4}$ | $p53^{i-i+7}$ | $p53^{double}$ |
|---|---|---|---|---|
| 1-2 | 0.35 | 0.35 | 0.34 | 0.35 |
| 2-3 | 0.35 | 0.36 | 0.36 | 0.36 |
| 3-4 | 0.36 | 0.36 | 0.36 | 0.36 |
| 4-5 | 0.36 | 0.36 | 0.36 | 0.36 |
| 5-6 | 0.36 | 0.36 | 0.36 | 0.36 |
| 6-7 | 0.36 | 0.36 | 0.36 | 0.36 |
| 7-8 | 0.37 | 0.37 | 0.36 | 0.36 |
| 8-9 | 0.37 | 0.37 | 0.36 | 0.36 |
| 9-10 | 0.36 | 0.37 | 0.36 | 0.36 |
| 10-11 | 0.37 | 0.37 | 0.37 | 0.37 |

#### 7. Free energy Surface comparison between Unbiased and T-REMD

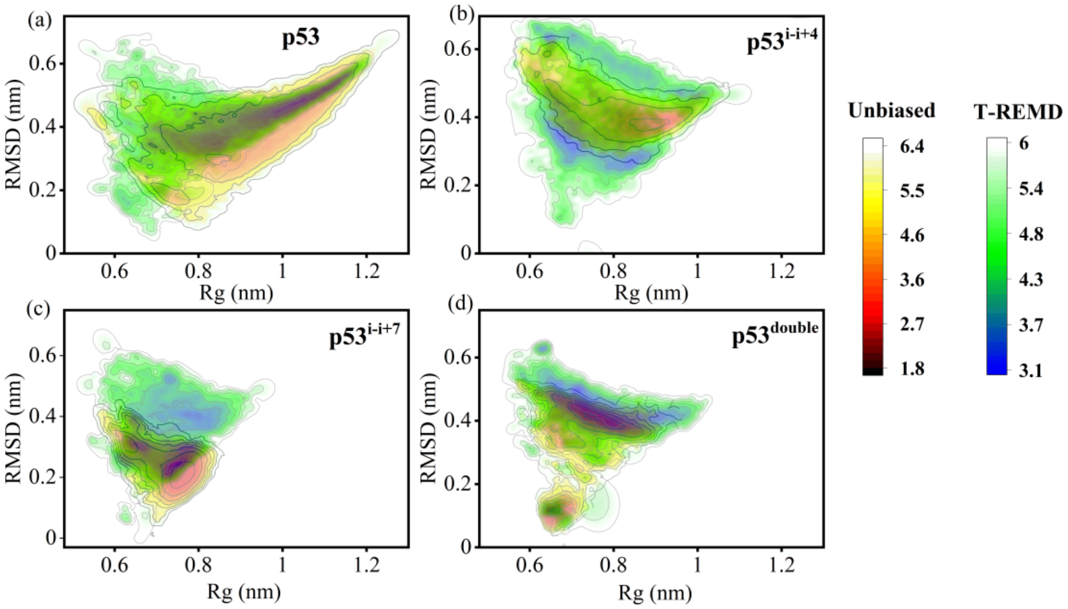


**Figure S6:** Comparison of Free energy surface along Rg and RMSD between unbiased and T-REMD simulation for systems (a) p53, (b) $p53^{i-i+4}$, (c) $p53^{i-i+7}$, and (d) $p53^{double}$ are represented with energy color bars. Free energy values are in $kcalmol^{-1}$.

#### 8. End-to-end Distance and Helical fraction free energy surfaces

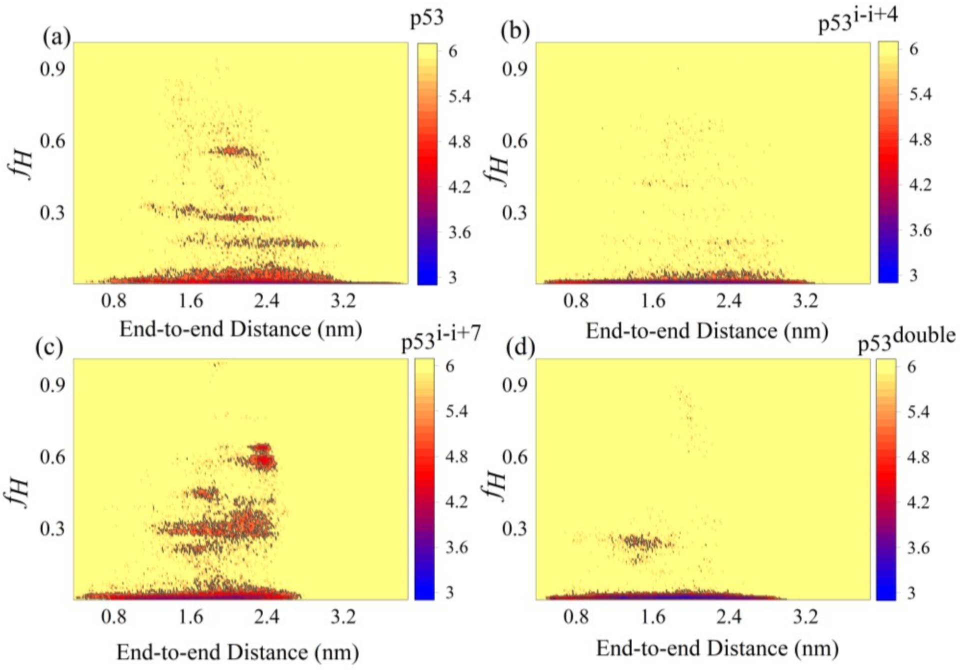


**Figure S7:** Free energy surface along end-to-end distance and helical fraction obtained from T-REMD simulations for systems (a) p53, (b) $p53^{i-i+4}$, (c) $p53^{i-i+7}$, and (d) $p53^{double}$ are represented with energy color bars. Free energy values are in $kcalmol^{-1}$.

#### 9. Helical fraction of peptide in MDM2 Bound state

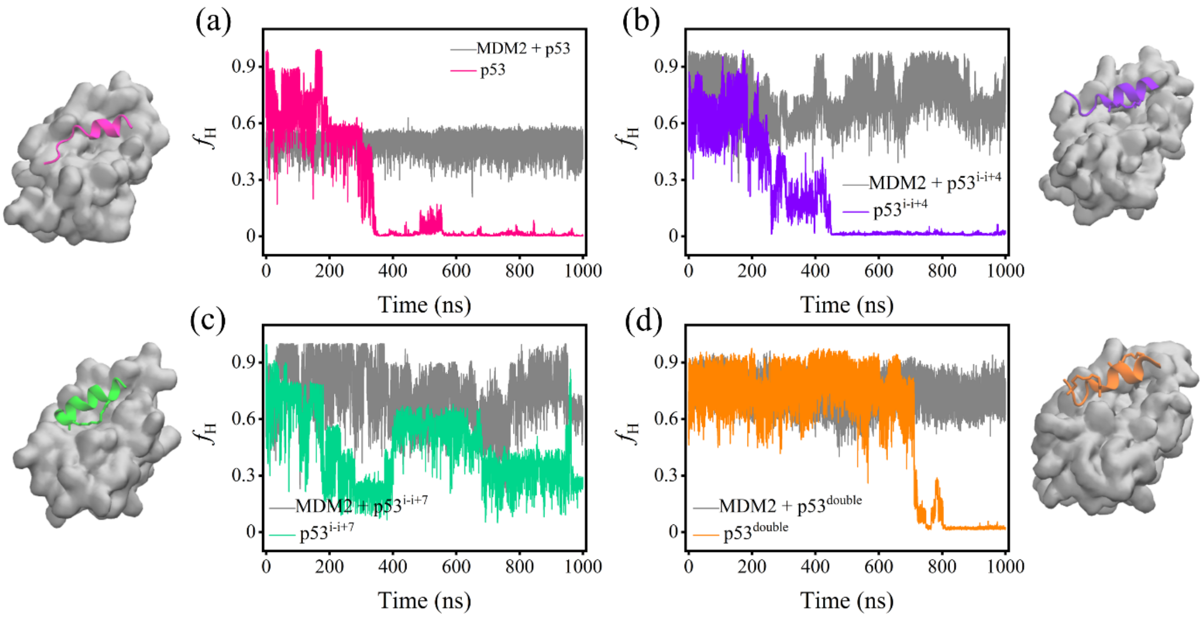


**Figure S8:** Time evolution of the helical fraction of p53 in MDM2 bound state and free state for (a) Wild-type p53, (b) $p53^{i\text{-}i+4}$, (c) $p53^{i\text{-}i+7}$, and (d) $p53^{double}$.

#### 10. Table ST2: Binding energy components 3 independent simulations of wild-type p53 with MDM2. All the energy components are in kcalmol$^{-1}$.

| Energy Components | Simulation1 | Simulation2 | Simulation3 | Average | Standard Deviation |
|---|---|---|---|---|---|
| ΔBOND | 1.15 | 9.42 | -0.1 | 3.49 | 4.22 |
| ΔANGLE | 4.35 | 1.19 | 1.24 | 2.26 | 1.48 |
| ΔDIHED | -3.99 | 6.32 | 2.12 | 1.48 | 4.23 |
| ΔUB | 0.01 | 0.32 | 0.01 | 0.11 | 0.15 |
| ΔIMP | 0.26 | 0.84 | 0.32 | 0.47 | 0.26 |
| ΔCMAP | 1.8 | 5.55 | 3.31 | 3.55 | 1.54 |
| ΔVDWAALS | -61.9 | -83.49 | -69.31 | -71.57 | 8.96 |
| ΔEEL | -257.41 | -365.47 | -261.27 | -294.72 | 50.05 |
| Δ1-4 VDW | -0.67 | 4.2 | 2.53 | 2.02 | 2.02 |
| Δ1-4 EEL | -15.64 | 13.75 | -2.6 | -1.50 | 12.02 |
| ΔEGB | 293.24 | 371.89 | 282.82 | 315.98 | 39.76 |
| ΔESURF | -9.51 | -11.22 | -9.36 | -10.03 | 0.84 |
| ΔGGAS | -332.03 | -407.37 | -323.76 | -354.39 | 37.62 |

| | | | | | |
|---|---|---|---|---|---|
| ΔGSOLV | 283.73 | 360.67 | 273.46 | 305.95 | 38.92 |
| ΔEnthalpy | -48.31 | -46.69 | -50.3 | -48.43 | 1.48 |

**11. Table ST3**: Binding energy components 3 independent simulations of p53$^{i\text{-}i+4}$ with MDM2. All the energy components are in kcalmol$^{-1}$.

| Energy Components | Simulation1 | Simulation2 | Simulation3 | Average | Standard Deviation |
|---|---|---|---|---|---|
| ΔBOND | -1.34 | -2.37 | -1.71 | -1.81 | 0.43 |
| ΔANGLE | -2.71 | -2.6 | -2.33 | -2.55 | 0.16 |
| ΔDIHED | 8.39 | 15.5 | 5.71 | 9.87 | 4.13 |
| ΔUB | -0.9 | -0.94 | -0.72 | -0.85 | 0.10 |
| ΔIMP | 0.22 | 0.23 | -0.43 | 0.01 | 0.31 |
| ΔCMAP | 7.15 | 1.21 | 6.83 | 5.06 | 2.73 |
| ΔVDWAALS | -76.88 | -78.73 | -74.18 | -76.60 | 1.87 |
| ΔEEL | -367.19 | -428.91 | -224.54 | -340.21 | 85.59 |
| Δ1-4 VDW | 4.2 | 3.45 | 3.68 | 3.78 | 0.31 |
| Δ1-4 EEL | 8.99 | 14.09 | 5.89 | 9.66 | 3.38 |
| ΔEGB | 363.85 | 412.78 | 236.98 | 337.87 | 74.08 |
| ΔESURF | -9.37 | -10.44 | -7.54 | -9.12 | 1.20 |
| ΔGGAS | -420.07 | -479.07 | -281.8 | -393.65 | 82.67 |
| ΔGSOLV | 354.48 | 402.34 | 229.44 | 328.75 | 72.89 |
| ΔEnthalpy | -65.59 | -76.73 | -52.36 | -64.89 | 9.96 |

**12. Table ST4**: Binding energy components 3 independent simulations of p53$^{i\text{-}i+7}$ with MDM2. All the energy components are in kcalmol$^{-1}$.

| Energy Components | Simulation1 | Simulation2 | Simulation3 | Average | Standard Deviation |
|---|---|---|---|---|---|
| ΔBOND | 8.33 | 7.91 | -0.07 | 5.39 | 3.86 |
| ΔANGLE | -0.89 | 1.13 | -5.08 | -1.61 | 2.59 |
| ΔDIHED | 6.51 | 2.94 | 7.64 | 5.70 | 2.00 |
| ΔUB | 0.25 | 0.49 | 0.05 | 0.26 | 0.18 |

| | | | | | |
|---|---|---|---|---|---|
| ΔIMP | 0.28 | 0.46 | -0.03 | 0.24 | 0.20 |
| ΔCMAP | 2.87 | 2.19 | 1.93 | 2.33 | 0.40 |
| ΔVDWAALS | -75.57 | -66.62 | -70.1 | -70.76 | 3.68 |
| ΔEEL | -249.66 | -234.59 | -277.38 | -253.88 | 17.72 |
| Δ1-4 VDW | 3.73 | 1.79 | 3.84 | 3.12 | 0.94 |
| Δ1-4 EEL | 22.49 | 9.91 | 17.36 | 16.59 | 5.16 |
| ΔEGB | 253.61 | 249.41 | 279.64 | 260.89 | 13.37 |
| ΔESURF | -9.48 | -8.7 | -8.97 | -9.05 | 0.32 |
| ΔGGAS | -281.66 | -274.39 | -321.83 | -292.63 | 20.86 |
| ΔGSOLV | 244.13 | 240.71 | 270.68 | 251.84 | 13.39 |
| ΔEnthalpy | -37.53 | -33.69 | -51.15 | -40.79 | 7.49 |

**13. Table ST5:** Binding energy components 3 independent simulations of $p53^{double}$ with MDM2. All the energy components are in $kcalmol^{-1}$.

| Energy Components | Simulation1 | Simulation2 | Simulation3 | Average | Standard Deviation |
|---|---|---|---|---|---|
| ΔBOND | -2.69 | -2.61 | -2.72 | -2.67 | 0.05 |
| ΔANGLE | -2.64 | -8.63 | -6.92 | -6.06 | 2.52 |
| ΔDIHED | 17.05 | 3.53 | 15.06 | 11.88 | 5.96 |
| ΔUB | -0.81 | -1.05 | -0.91 | -0.92 | 0.10 |
| ΔIMP | -0.2 | 0 | 0.26 | 0.02 | 0.19 |
| ΔCMAP | -5.48 | 9.58 | 1.1 | 1.73 | 6.16 |
| ΔVDWAALS | -68.13 | -74.01 | -72.92 | -71.69 | 2.55 |
| ΔEEL | -269.76 | -237.14 | -259.92 | -255.61 | 13.66 |
| Δ1-4 VDW | 0.81 | 4.19 | 3.18 | 2.73 | 1.42 |
| Δ1-4 EEL | -17 | 14.59 | -2.72 | -1.71 | 12.92 |
| ΔEGB | 287.01 | 229.49 | 267.04 | 261.18 | 23.85 |
| ΔESURF | -9.31 | -9.03 | -8.64 | -8.99 | 0.27 |
| ΔGGAS | -348.85 | -291.54 | -326.51 | -322.30 | 23.59 |
| ΔGSOLV | 277.7 | 220.46 | 258.4 | 252.19 | 23.78 |
| ΔEnthalpy | -71.15 | -71.08 | -68.11 | -70.11 | 1.42 |

#### 14. Hydrogen bonds between MDM2 and stapled p53:

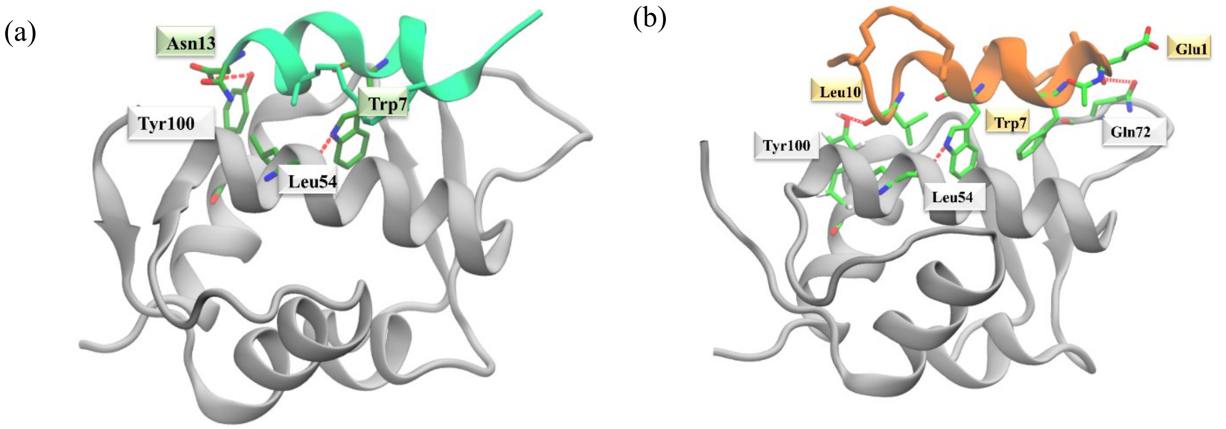


**Figure S9:** The relative positions of key residues showing hydrogen bond interactions with MDM2 are shown for (a) MDM2 + p53$^{i\text{-}i+7}$ and (b) MDM2 + p53$^{double}$. Hydrogen bonds are represented using red dashed lines.
